# *Resilin,* the gene for the molecular spring: its roles in flight and jumping of *Drosophila*

**DOI:** 10.1101/2025.10.01.678482

**Authors:** Miyuna Hagiwara, Yoshiki Eto, Yoko Keira, Hiroyuki O. Ishikawa, Takaomi Sakai, Toshiro Aigaki, Tsunaki Asano

## Abstract

Resilin is a member of the chitin-binding protein family that was originally found as a major component of an elastic matrix present in insects. The knockdown of the gene for this protein in *Drosophila melanogaster* causes the characteristic downturned wing posture, and the knockdown flies cannot move their wings. The jump distance of the knockdown flies is around 50% shorter than that of the controls. Null mutant flies show the same phenotypes, which can be rescued by introducing a 4.7 kb genomic fragment harboring the whole coding region of *Resilin*. *In vitro* experiments have shown that the elasticity of the matrix made of Resilin is due to the dityrosine-mediated polymerization of Resilin molecules. Dual oxidase (Duox) is the most likely candidate for the *in vivo* polymerization of Resilin molecules. *Duox* knockdown induces phenotypes similar to those observed in wing posture and jump performance of the *Resilin* knockdown and knockout flies, which can be rescued by the overexpression of *Duox* gene from a beetle. These findings suggest that Duox is an essential factor for the proper function of Resilin as the major component of the resilin matrix.

## Introduction

Insects are often regarded as the most successful animals on Earth with their extreme diversity, enormous biomass, and great impact on the global ecosystem. There have been intensive debates as to why insects have been so successful, and their ability to fly must have had great impacts on their evolutionary process and the expansion of their habitat to a wide range of terrestrial environments. It is considered that the ability to fly is advantageous in the competition for survival and the creation of new niches not easily accessible to other animals, which is supported by the fact that 99% of the described insect species are winged.

During powered flight, insects flap their wings by moving their flight muscles in the thorax. Repetitive and rapid contractions of flight muscles are transformed into high-frequency wing motion, generating a force to lift the insect’s body^1,2^. The hinges connecting the wings and the body have thick tendons, which may be involved in energy transformation from the flight muscles to the wings^3,4^. Accumulation of mechanical loads derived from the high-frequency vibration during flight could eventually destroy structures around the wing hinges. However, the thick tendon of the wing hinges is composed of a rubber-like matrix, which may presumably absorb/relieve such stresses, and prevent the hinges from breaking down^3,4,5^. In addition to their powered flight, insects may also use the properties of such rubber-like matrices for jumping. Similarly to flight, jumping is a high-velocity locomotion through the air, which may be another advantage of insects that is not found in other invertebrates, with only a few exceptions of some arthropods, such as sand hoppers or jumping spiders^6,7,8^. It has been reported that the rubber-like matrices are found in the joints of the hind legs of the locust or in the pleural arch at the top of the hind legs of the flea. Both of which are positioned in places that can be easily imagined to be used as a spring providing highly efficient energy kinetics for achieving long-distance and high-velocity locomotion^7,9,10^.

The first description of the rubber-like matrices in insects was the discovery of tendons in the wing hinges of dragonflies and locusts^4^. At that time, the rubber-like matrices were named “resilin” after the Latin word “resilire,” which means resilience, and were defined as materials that can be stained with methylene blue or toluidine blue. After that, rubber-like matrices (hereafter, referred to as the “resilin matrix”) were found in the salivary pump and feeding pump of bags, or the tymbal of cicadas for sound production^11,12^. The connection between jumping mechanisms and the localization of the resilin matrix at leg joints has also been discussed in insects such as locusts that have the thick resilin matrix at the joints of their hind legs^9,13^. Relatively recent reports on the functions of the resilin matrix include examples from bed bugs and midges. Male common bedbugs (*Cimex lectularius* and *Cimex hemipterus*) puncture the intersegmental membrane of the female abdomen with a needle to inject their sperm. After the male detaches from the female, the pore is likely closed, probably owing to the elasticity of the resilin matrix^14,15^. Larvae of the phantom midge (*Chaoborus americanus*) can regulate their buoyancy by altering the volume of their air sacs. This is achieved through a pH-dependent transition in the shape of the resilin matrix that composes the air sacs. During this process, membrane ATPase controls the proton content of the resilin matrix^16^.

The physical properties of the resilin matrix are not affected considerably by deep freezing, high temperature, or treatments with alcohols or fixatives. In contrast, the resilin matrix is rapidly dissolved by proteolytic digestion, indicating that proteins are its major constituents^4^. Subsequently, a study showed that proteins in the resilin matrix are crosslinked together through the formation of di-or tri-tyrosine, in which tyrosine residues are covalently crosslinked to each other^17^. It was also shown that di- and tri-tyrosine emit the characteristic blue fluorescence under UV irradiation, and this property has been used as a useful tool to observe the localization of the resilin matrix noninvasively (hereafter, di- and tri-tyrosine are collectively referred to as dityrosine). In 2001, Ardell and Andersen reported a candidate gene encoding a protein composing the resilin matrix. Partial amino acid sequences of peptides isolated from the protease digest of the resilin matrix taken from the desert locust *Schistocerca gregaria* were determined by Edman degradation. The obtained sequences were used for BLAST search in the genomic database of the fruit fly *Drosophila melanogaster* to find candidate genes encoding *Drosophila* proteins homologous to the locust proteins^18^. Two proteins (CG15920 and CG9036) were found to have the sequences that matched well with those obtained from the resilin matrix of the locust. On the basis of similarity in amino acid compositions (high amounts of proline, glycine, and tyrosine), Ardell & Andersen concluded that CG15920 was the most likely candidate, and tentatively named the *CG15920 “Resilin*” as the gene for a major protein composing the resilin matrix. Since that study, *CG15920* has been called *Resilin* (hereafter, *Rsl*). Until recently, however, there had been no reports on the gene function analysis to directly demonstrate that the gene encoding resilin is actually involved in insect flight or jumping. Furthermore, it was also not known how resilin molecules are polymerized *in vivo*, although *in vitro* experiments using a recombinant Rsl protein of *D. melanogaster* have shown that Rsl polymerization is necessary for the high elasticity of synthetic resilin matrix^5,19–21^.

Here, using the newly obtained *Rsl* mutant of *D. melanogaster*, we characterized the phenotypes and showed that *Rsl* has roles in preventing the breakdown of the wing hinge structure over time and in efficient jumping. We also tried to show which genes were involved in the polymerization of the Rsl molecule. Dual oxidase (Duox) is a membrane protein with the extracellular peroxidase domain and the transmembrane NADPH-oxidase domain. By tissue-specific *Duox* knockdown and its rescue experiment, we showed that Duox was indeed the factor for Rsl polymerization. We also showed that the Duox-mediated polymerization of Rsl was indispensable for the proper function of the Rsl/resilin matrix both in flight and jumping, which may be the critical factors for the evolution and success of insects as terrestrial organisms.

## Results

### *Rsl* encodes a protein with “elastic repeats”

*Rsl* is composed of three exons, and two isoforms *Rsl-RA* and *Rsl-RB* (with and without exon 2, respectively) are expressed (Figure 1A)^22,23^. Rsl is the secreted protein with a signal sequence (Figure 1B)^18^. The polypeptide encoded by *Rsl-RA* is divided into three parts: 1) repeats of N-terminally located “elastic modulus”, 2) chitin-binding motif, and 3) repeats of C-terminally located “elastic modulus”. The polypeptide encoded by *Rsl-RB* is devoid of the large portion of the chitin-binding motif (Figure 1B)^18,24^.

**Figure 1.**
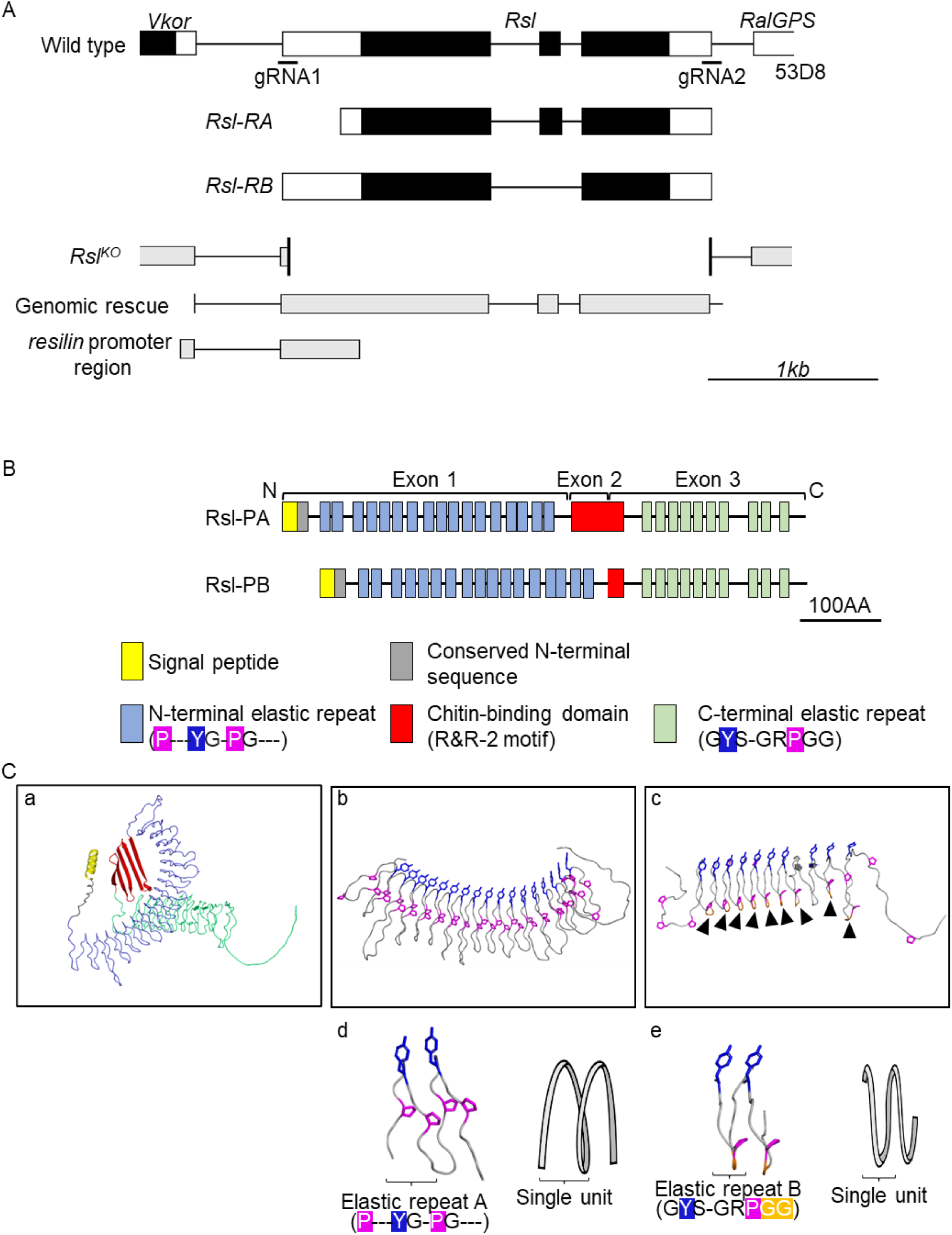
Exon–intron organization in the *Rsl* locus and protein structures of Rsl. (A) The top is a schematic of the *Rsl* locus in the wild-type fly. Black and white boxes are CDSs and UTRs, respectively. The exon compositions of the two isoforms are shown below. The schematics below show the *Rsl* locus in *Rsl^KO^* flies; the regions chosen to generate the transgenes for the rescue experiments (*Rsl-GR*) and for the Gal4 driver (*Rsl-Gal4*) are shown. The short bars indicate two gRNAs designed for the deletion of CDS to generate *Rsl^KO^* flies. (B) Domain-motif organization of Rsl isoforms are shown. Each of the functional domain or consensus is shown below. Rsl-PB has a small portion of the chitin-binding R&R consensus sequence (red box) encoded by the 5’ part of exon 3. (C) The 3D model of Rsl-PA predicted in AlphaFold Server is shown. Panel a is the whole Rsl-PA model. Blue and green parts are the N-terminal and C-terminal repeats, respectively. In panels b and c, the N-terminal and C-terminal repeats are enlarged with their proline and tyrosine residues illustrated by blue and purple colors, respectively. In panel c, beta turns are indicated by closed arrowheads. Panels d and e show the views in which two repeats are extracted from panels b and c, respectively, with schematics for two cycles of their polypeptide backbones.

The elastic moduli^24^ (hereinafter referred to as elastic repeats) that are indicated by light-blue or light-green boxes (Figure 1B) are characterized by the consensus sequences containing proline, glycine, and tyrosine. The consensus sequences are also found in other insect proteins that are annotated as resilin^23,24^. The tyrosine residue is considered important for the formation of intermolecular crosslinks^18^. As first described by Ardell and Andersen^18^, the consensus sequences of the elastic repeats in the N- and C-terminal regions are different from each other (Figure 1B). In *D. melanogaster*, the N-terminal repeats (-**P**--<u>Y</u>G-**P**G---) have two prolines with an intervening tyrosine, whereas the C-terminal repeats (G<u>Y</u>S-GR**P**GG) have only one tyrosine and one proline. In the multiple sequence alignment^23,24^, the N-terminal repeats are conserved among species, but there are at least two subtypes in the consensus sequences of the C-terminal repeats. In many insects, the consensus is G<u>Y</u>**P**SGG**P**GG with two prolines (type-A repeat). In contrast, many fly species and a portion of hemipteran insects have different consensus sequences characterized by the presence of only one proline (Figure 1B). In a comprehensive database search of arthropod genomes, we found that in amino acid sequences of resilins from the palaeopteran insects (Odonata and Ephemeroptera: the orders of the most primitive winged insects) or those from the collembolan (the primitive non-insect hexapods), no consensus sequences of the C-terminal repeats are found (Figure S1). Considering the branching pattern of the phylogenetic tree illustrating the evolutionary process of insects (see Figure S1), it can be hypothesized that the C-terminal repeats are a trait that was obtained in accordance with the emergence of a new type of winged insects (Neoptera). In the three-dimensional model of *Drosophila* Rsl (Q9V7U0 in the uniplot database), which was predicted using AlphaFold Server (https://alphafoldserver.com) powered by AlphaFold 3^25^ (Figure 1C), both the N- and C-terminal regions have a spring-like “coiled” shape (panels b and c, respectively), with each elastic repeat corresponding to one cycle of the spring (panels d and e). The presence of the beta turn at the short PGG sequence motif in the C-terminal repeats (arrowheads in panel c) is consistent with the previous prediction^18^. Our predictions using AlphaFold 3 showed a spring-like shape similar to the Rsl model in the resilin of other insects, except for species such as *Bombyx mori* (Lepidoptera) and *Folsomia candida* (Collembola) (Figure S2). The spring-like “coiled” shape in the models is intriguing, considering the function of the resilin as a component of spring-rubber-like biomaterials.

### *Rsl* is expressed in the wing hinge and the leg joint

In previous studies of the two Drosophilidae species^22,26^, it was shown that *Rsl* was mainly expressed in late pupal stages. In our RT-PCR analysis (Figure 2A), the apparent signals of the isoforms *Rsl-RA* and *Rsl-RB* were detected from the pupal to early adult stages. From 2 to 4 days after puparium formation (APF), the signals for *Rsl* expressions are strong, and the emergence of the signal for the expression of *Rsl-RB* precedes that of *Rsl-RA*. In western blotting with the specific antibody, two signals corresponding to the molecular masses of 57k and 52k can be seen (Figure 2B). The upper and lower signals correspond to mature Rsl-PA and Rsl-PB without a signal sequence, respectively (Figure S3). The emergence of the Rsl-PB signal precedes that of Rsl-PA, which is consistent with the temporal patterns obtained by RT-PCR analysis (Figure 2A). Rsl signals almost disappear after eclosion, which is consistent with the observation that resilin molecules are polymerized to become insoluble materials^17^. Rsl polymer cannot penetrate into the SDS-PAGE gel and are not detectable by western blot analyses.

**Figure 2.**
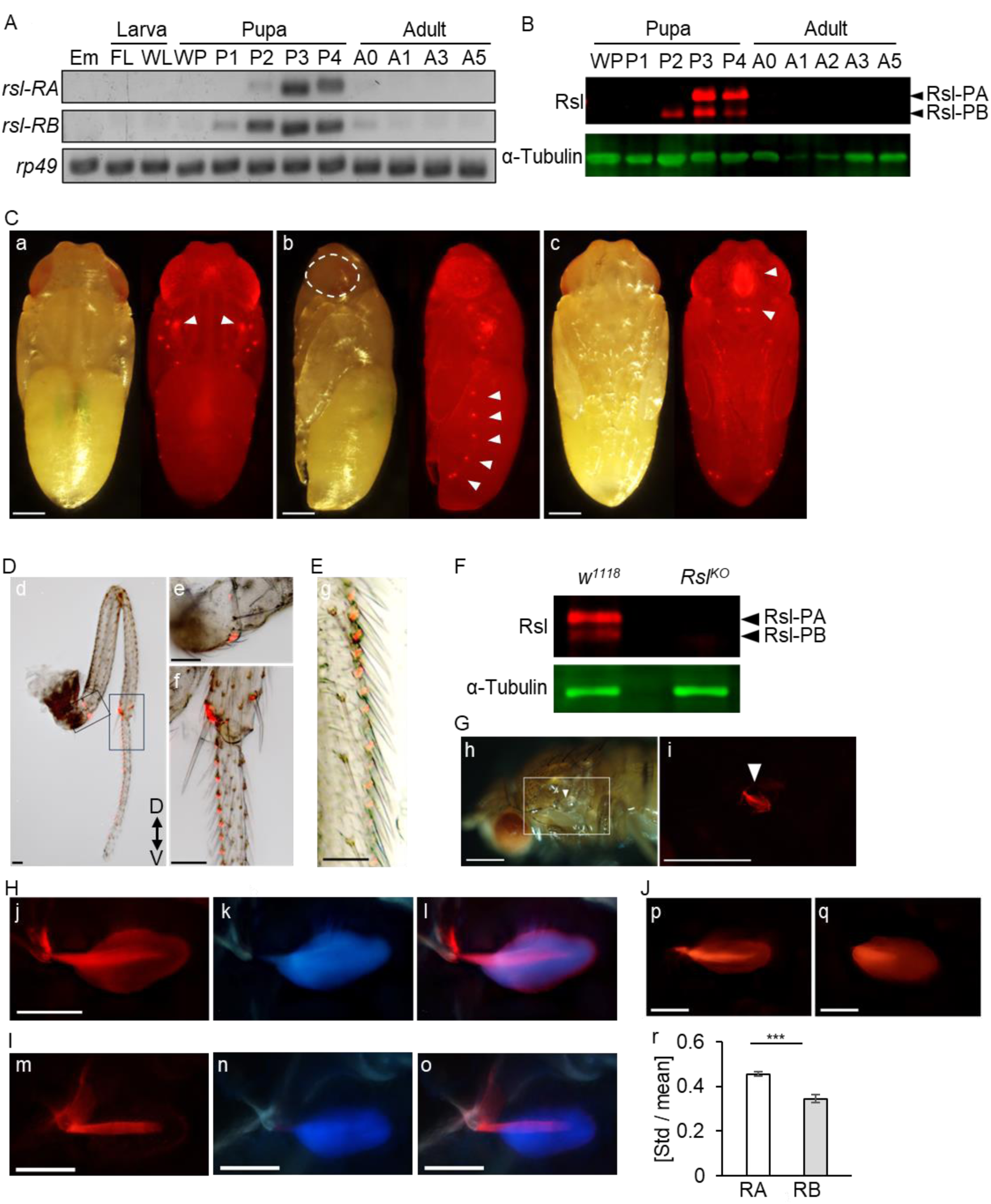
Temporal patterns of *Rsl* expression and spatial patterns of the reporter. (A) RT-PCRs of *Rsl-RA* and *Rsl-RB* from embryo to day 5 adult are shown. (B) Western blots of the Rsl protein from white pupa to day 5 adult are shown. In both A and B, FL, WL, WP, P1-4, and A0-5 are the abbreviations of feeding larva, wandering larva, white pupa, pupa days 1–4, and adult day 0–5, respectively. (C–H and J) Spatial distribution of the Rsl-RA::mCherry reporter expressed with *Rsl-Gal4* is shown. (C) Panels a – c show the views from dorsal, lateral, and ventral directions, respectively. Photos on the left side were taken under white light. In each panel, white arrowheads indicate the mCherry signals at the wing hinges, spiracles, or labrum. The dotted circle indicates the outline of the compound eye. (D and E) The midleg of the flies (D) and the edge of the wing (E) are shown. Photos of the mCherry signal and bright field were merged. Scale bar: 50 µm. (F) The western blots of Rsl in the compound eyes dissected from day-3 pupae of the wild-type and *Rsl^KO^*flies are shown. The electrophoretic mobilities of Rsl-PA and Rsl-PB are indicated with arrowheads. (G) The Rsl-RA::mCherry localization at the wing hinge is shown. The outer cuticle was removed together with the wing (arrowhead) to observe the internal structure (panel h). In panel i, the area enclosed in the square in panel h is enlarged. Scale bar: 500 µm. (H) Panel j is the magnified view of the wing hinge of the fly expressing *Rsl-RA::mCherry*. Panel k is the photo taken under UV radiation, and panel l is the merged photo of panels j and k. Scale bar: 50 µm. (I) The distribution pattern of ChtVis-tdTomato that was expressed with *Rsl-Gal4* is shown. Scale bar: 50 µm. In both H and I, the wavelengths of the filters used for excitation and the detection of dityrosine are 330–385 and >420 nm, respectively. (J) The biased distributions of Rsl::mCherry signals are compared. Panels p and q are the photos of the resilin matrix visualized with Rsl-RA::mCherry and Rsl-RB::mCherry, respectively. In panel r, the deviations of the biased distribution were calculated as described in Materials and Methods.

To see the spatial patterns of *Rsl* expression, we prepared the *Rsl* promoter-Gal4 driver that covers the genomic region around 2.3 kbp upstream of the first ATG (Figure 1A). The expression was visualized using the *UAS*-*Rsl-RA*::*mCherry* reporter. The photos in Figure 2C show pupae on day 3 APF. The mCherry signal was visible at the hinges of the wings, spiracles, compound eyes, and labrum (Figure 2C, panels a, b, and c). In the observation of the adult on day 3 after eclosion, the mCherry signal can be seen on the ventral side of the leg joints between the trochanter and the femur, and the roots of the mechanosensory hairs of the leg and wing (Figures 2D and 2E). These expression patterns were obtained using flies carrying the *Rsl-*promoter Gal4 sequence integrated into the 3rd chromosome. The mCherry signal pattern obtained with the *Rsl*-promoter Gal4 sequence on the second chromosome was nearly identical to the data obtained with *Rsl-*promoter Gal4 on the third chromosome, except that the signal in the compound eyes was very weak (Figure S4). In western blot analysis of the compound eye (Figure 2F), the signals of Rsl can be seen in the wild-type fly, and this signal disappeared in the sample from *Rsl^KO^* that we generated (Figures 1A and S5). The results of the western blot analysis indicate that the signal in the wild-type fly corresponds to Rsl, and that the compound eye surely expresses *Rsl*.

To observe the details of the internal structure at the wing hinge, we removed the outer cuticle. The mCherry signal was clearly visible just beneath the wing hinge (Figure 2G, panel i), and the structure exhibited blue fluorescence under UV irradiation (Figure 2H). The intensity of this blue fluorescence signal decreased markedly and increased under acidic and alkaline conditions, respectively (see Figure S6), consistent with the characteristics of dityrosine previously reported^17^. This structure, which is positive for both the mCherry signal and blue fluorescence, is the tergopleural tendon. Previous studies have shown that this tendon is the structure connecting the direct flight muscle and the root of the wing at the wing hinge^27–29^.

Since Rsl-PA has the chitin-binding domain (Figure 1B), we consider that this domain may function in chitin binding. To visualize the distribution of the chitin fiber inside the tergopleural tendon, we expressed ChtVis-tdTomato with *Rsl-*promoter-Gal4. A strong signal can be seen along the horizontal midline, indicating the presence of chitin along the midline (Figure 2I). This is similar to the spatial pattern of Rsl-RA::mCherry (Figure 2H). Next, we compared the distributions of Rsl-RA::mCherry and Rsl-RB::mCherry (with and without the sequence of the chitin-binding domain, respectively) that were separately expressed (Figure 2J). The spatial pattern of Rsl-RA::mCherry, which is denser along the horizontal central line (panel p), contrasts with the more uniform distribution of Rsl-RB::mCherry (panel q). In the calculation for the biased distribution of mCherry signals (see Materials and Methods), the value was higher in flies expressing Rsl-RA::mCherry than in those expressing Rsl-RB::mCherry. This indicates a more uniform distribution of Rsl-RB::mCherry, which may be caused by the absence of a chitin-binding domain.

### *Rsl* may work to prevent the breakdown of a tendon in the wing hinge that is required for sustainable flight

To examine the effects of the suppression of *Rsl* expression, we knocked down *Rsl*^30^ with the *Rsl*-promoter Gal4 driver, and the expression was suppressed to around the baseline levels (Figure S7). In the *Rsl* knockdown flies, no visible phenotypes were observed in the very early stages of the adult, but after one day, all the knockdown individuals were unable to move their wings and started to display the characteristic downturned wing posture (Figure 3A). The same wing posture has been observed in multiple studies including the analysis of *pink1*, which functions in muscle development^31,32^. The phenotypes observed in our *Rsl* knockdown are the same as those previously reported for the *Rsl* knockdown flies^21^. To further characterize the function of *Rsl*, we used the null mutant (*Rsl^KO^*) that was prepared in this study (Figure S5). The homozygous *Rsl^KO^* flies showed the characteristic downturned wing posture, and they could not move their wings, which are the same phenotypes as those observed in *Rsl* knockdown flies (Figure 3C, see Extended data Video 2). To rescue these phenotypes, we generated flies with the transgene of the 4.7 kbp genomic fragment harboring the *Rsl* coding region on the 3rd chromosome (*Rsl*-Genomic rescue: *Rsl-GR*) (Figure 1A). The *Rsl^KO^* flies with *Rsl-GR* did not show the downturned wing posture (Figures 3C and 3D), and they could move their wings and fly (see Supplementary mov. S3), indicating that *Rsl-GR* rescued the phenotypes of *Rsl^KO^*. Then, we also tried to rescue the phenotypes of *Rsl^KO^* using the Gal4/UAS system (Figure 3D). Although flies expressing each isoform only gradually showed the wing posture phenotype, the phenotype was almost rescued. Combining the two isoforms did not show markedly different results with those obtained from either *Rsl-RA* or *Rsl-RB* alone.

**Figure 3.**
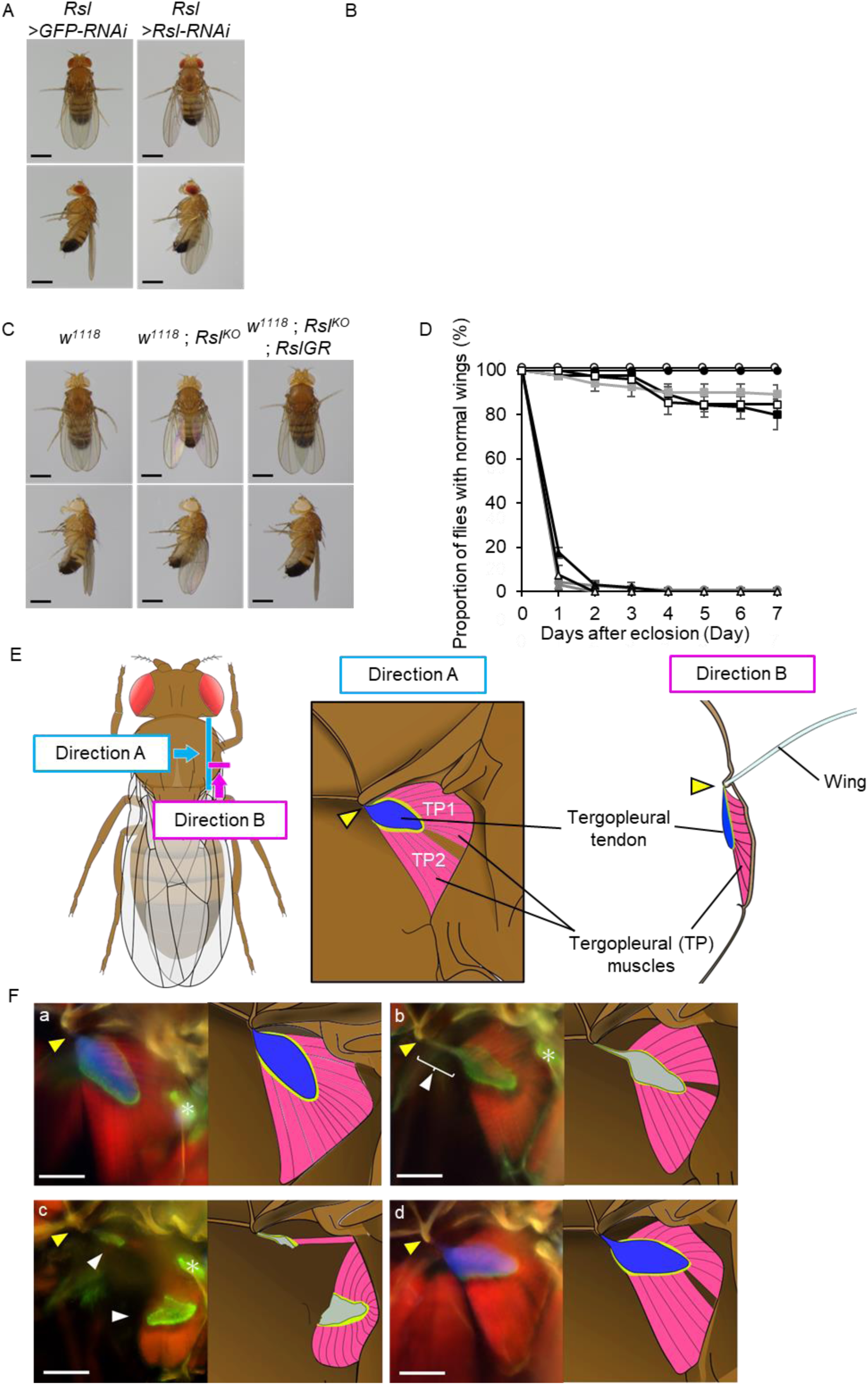
Phenotypes of *Rsl* knockdown and mutant flies. (A) The wing postures of the control fly (left: *Rsl>GFP-RNAi*) and *resilin* knockdown fly (right: *Rsl>Rsl-RNAi*) are compared. Scale bar: 0.5 mm. (B) The proportion of flies with normal wing posture on each day against the total number of flies is indicated (closed circles: *Rsl>GFP-RNAi*, grey circles: *Rsl>Rsl-RNAi*). (C) The wing postures of wild-type (*w^1118^*)*, Rsl^KO^* (*w^1118^*; *Rsl^KO^*), and rescued (*w^1118^*; *resilin^KO^*; *resilinGR*) flies are compared. Scale bar: 0.5 mm. (D) The temporal changes in the proportion of flies with normal wing posture after eclosion are shown. The fly genotypes are *w^1118^* (closed circles), *resilin^KO^* (gray circles), *resilin^KO^*; *resilinGR* (open circles), *resilin^KO^*; *resilin-Gal4* (closed triangles), *resilin^KO^, UAS-resilin-RA* (gray triangles), *resilin^KO^, UAS-resilin-RB* (open triangles), *resilin^KO^, UAS-resilin-RA; resilin-Gal4* (closed squares), *resilin^KO^,UAS-resilin-RB; resilin-Gal4* (gray squares), and *Rsl^KO^,UAS-Rsl-RA/Rsl^KO^,UAS-Rsl-RB*; *Rsl-Gal4* (open squares). (E) The middle (Direction A) is a schematic diagram of the wing hinge that is viewed from the midline toward the wing (blue arrow in the dorsal view of the fly). The right (Direction B) shows the wing hinge viewed from posterior to anterior direction (red arrow in the dorsal view of the fly). Blue shows the tergopleural tendon, which connects to the cuticle where the wing attaches (yellow arrowheads). Red indicates direct flight muscles TP1 and TP2^52^. (F) The inner structure of the wing hinge in *Dpy-YFP* was observed (panel a, left part), and its schematic is shown (right part). The red signals indicate muscles stained with phalloidin. The blue signal corresponds to blue fluorescence emitted upon UV irradiation. Green signals indicate Dpy-YFP. White arrow heads indicate the cuticle structure beneath the wing hinge attached to the proximal end of the tergopleural tendon. Panels b and c show the inner structures and the illustrations of the wing hinge of *Rsl^KO^* flies (with and without the downturned wing phenotype, respectively). Panel d shows those of the *Rsl^KO^* fly with *Rsl-*GR. In Figure 3F, the adults 72 h after eclosion were used for taking the photos, except for panel b for which adults 0 h after eclosion were used. Yellow arrowheads indicate the internal cuticle structure corresponding to the root of the wing hinge. White arrowheads indicate the tergopleural tendon devoid of any blue fluorescence signal in *Rsl^KO^* flies. Asterisks indicate Dpy-YFP signals attached to flight muscles other than TP-1 and TP-2. Scale bar: 50 µm.

It was shown in *D. melanogaster* that the tergopleural tendon was stained with the anti-Rsl antibody^28^, which is consistent with the presence of the signal for the Rsl::mCherry reporter at the tergopleural tendon (Figures 2G and 2H). In Figure 3E, the structure around the tergopleural tendon of the wild-type fly was illustrated according to the previous descriptions ^28,29^. The proximal side of the tergopleural tendon is attached to the inner side of the cuticle located at the wing hinge (indicated by the yellow arrowhead), and the direct flight muscles (tergopleural muscles 1 and 2: TP-1 and TP-2, respectively) are attached to both the tergopleural tendon and the cuticle of the body wall. In observation of the inner structure of wing hinges, *dumpy*-*YFP* (*dpy-YFP*) was used to visualize the boundary between the tendon and the muscle (Figure 3F, panel a). The green signal of Dpy-YFP surrounds the tergopleural tendon emitting the blue fluorescence upon UV irradiation (Figure 3F, panel a). Like the schematic figure (Figure 3E), the direct flight muscles are attached to the tergopleural tendon that is visualized with Dpy-YFP. As shown in panel b of Figure 3F, in the newly emerged *Rsl^KO^* fly, no blue fluorescence signal of dityrosine was seen. The loss of dityrosine signals can be explained by the absence of Rsl, which is supposed to provide the tyrosine residues as the source of dityrosine. The tergopleural tendon of the newly emerged *Rsl^KO^*fly remained linked to the root of the wing (white arrowheads), which may correlate with the absence of wing posture or movement abnormalities (Figure 3D). In contrast, in *Rsl^KO^* flies exhibiting the downturned wing posture (panel c), the tergopleural tendon was ruptured into two pieces (white arrowheads), confirming that the connection between the muscle and the root of the wing was almost lost. In contrast, in *Rsl^KO^* flies with *Rsl-GR* (panel d), the blue fluorescence that is surrounded by the Dpy-YFP signal is clearly visible. The structures of TP-1 and TP-2 are the same as those observed in wild-type flies, and no damage was observed. This is consistent with the *Rsl^KO^* flies with *Rsl-GR* showing no phenotypes of wing posture and movement abnormalities (Figures 3C and 3D; Extended data Video 3).

In the fields of anatomy and biomechanics, many reports define “resilin” as an extracellular structure that exhibits blue fluorescence, which is characteristic of dityrosine^33^. The blue fluorescence signal has been found in various body parts, including the intersegmental membrane, compound eyes, and wings^34,35^. In this study, to characterize the nature of the blue fluorescence in detail, we focused on the wings, since the presence of the blue fluorescence signal in wings has been reported in multiple insect species^34,35^. In the fly with *Rsl-RA::mCherry* expression that was driven by *Rsl-Gal4*, (Figure S8, panels, a and b), strong mCherry signal can be seen in several places including the tergopleural tendon. In contrast, there is almost no signal at the wing, although with UV, blue fluorescence can be seen at the proximal part of the wings (panel c). This finding suggests that *Rsl* is not expressed even in the area positive for blue fluorescence. In the magnified photo of the wild type fly’s wing (Figure S8, panel e), the arrows indicate distinct blue fluorescence signals at the proximal part of the wing, roughly corresponding to the area with landmarks adopted in a previous study (panel d)^36^. These signals became weaker drastically under acidic conditions (panel f), suggesting that they corresponded to dityrosine. In the wing of the *Rsl^KO^* fly (panel g), however, the signals were still seen, indicating that not Rsl but other tyrosine-rich substances are the cause of the blue fluorescence signal. These findings support the idea that dityrosine-rich structures exist without *Rsl* in *D. melanogaster*.

### Rsl is polymerized by the enzyme for ROS production

It has been shown that Duox is involved in dityrosine formation in ecdysozoans such as nematodes and insects^37–39^. The ROSs produced in the peroxidase domain may oxidize a tyrosine residue to form a tyrosine phenoxyl radical, followed by the reaction with the neighboring tyrosine residue to produce dityrosine or trityrosine^37,38^. Duox may be the most plausible candidate factor responsible for Rsl polymerization, because Rsl molecules are polymerized through dityrosine formation^17^. To demonstrate this, we examined the effect of *Duox* knockdown on Rsl polymerization or other phenotypic characteristics. In this study, we performed tissue-specific *Duox* knockdown with *Rsl-Gal4*, because whole body knockdown or knockout of *Duox* leads to lethality^37,40^. We used two UAS-lines, NIG stock #3131R-3 (*NIG3131R-3*) and the previously reported *UAS-Duox-RNAi_976-1145_*^40^, for *Duox* knockdown. In both cases, immediately after eclosion, *Duox* knockdown flies were able to fold and move their wings normally, but within a day, all of them exhibited the downturned wing posture, similar to *Rsl* knockdown and knockout (Figures 4A and 4B). After emergence of this wing posture, both the *Duox* knockdown flies could not move their wings, similarly to *Rsl* knockdown and knockout flies (hereafter, we mainly performed phenotypic characterization using *NIG3131R-3*).

**Figure 4.**
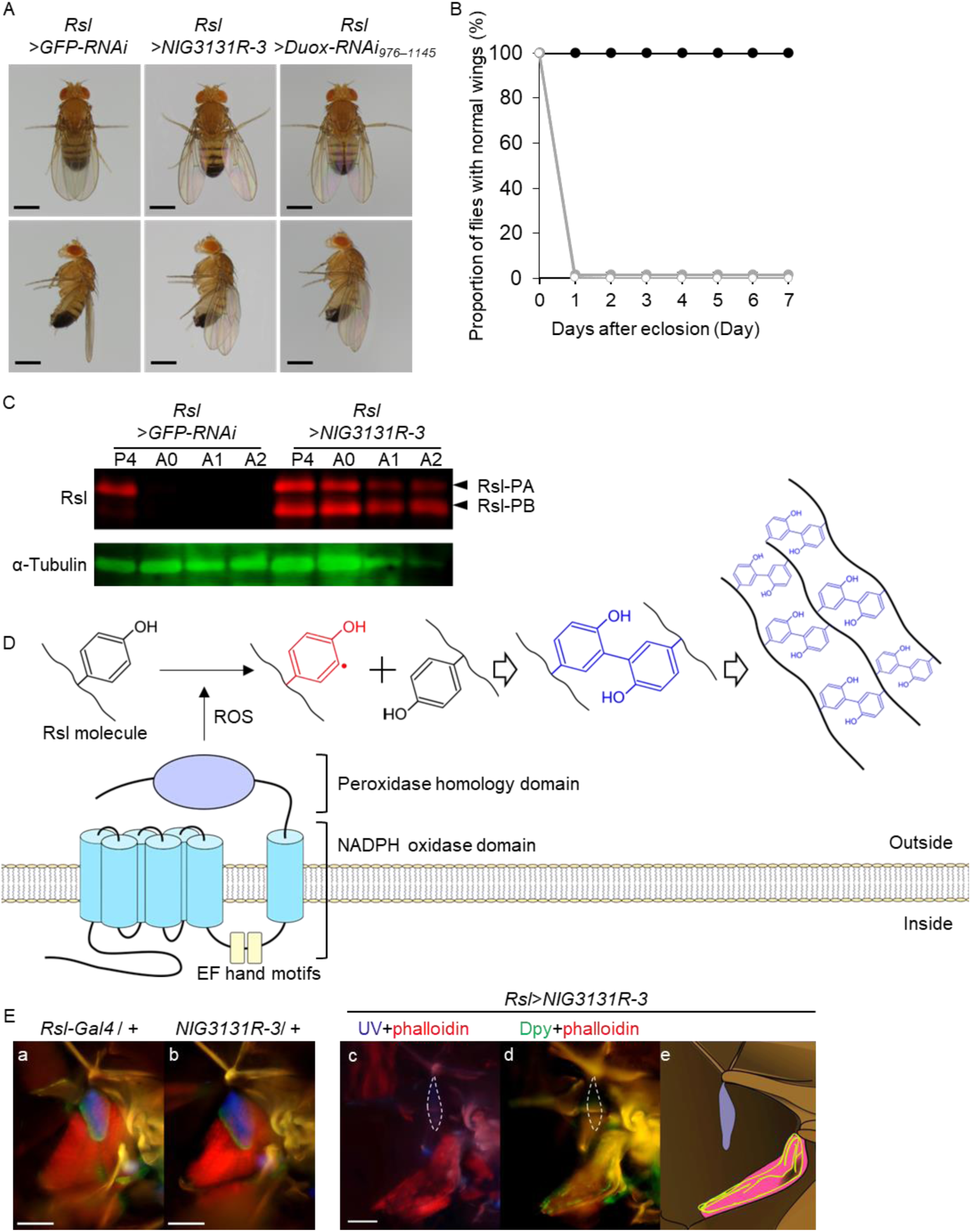
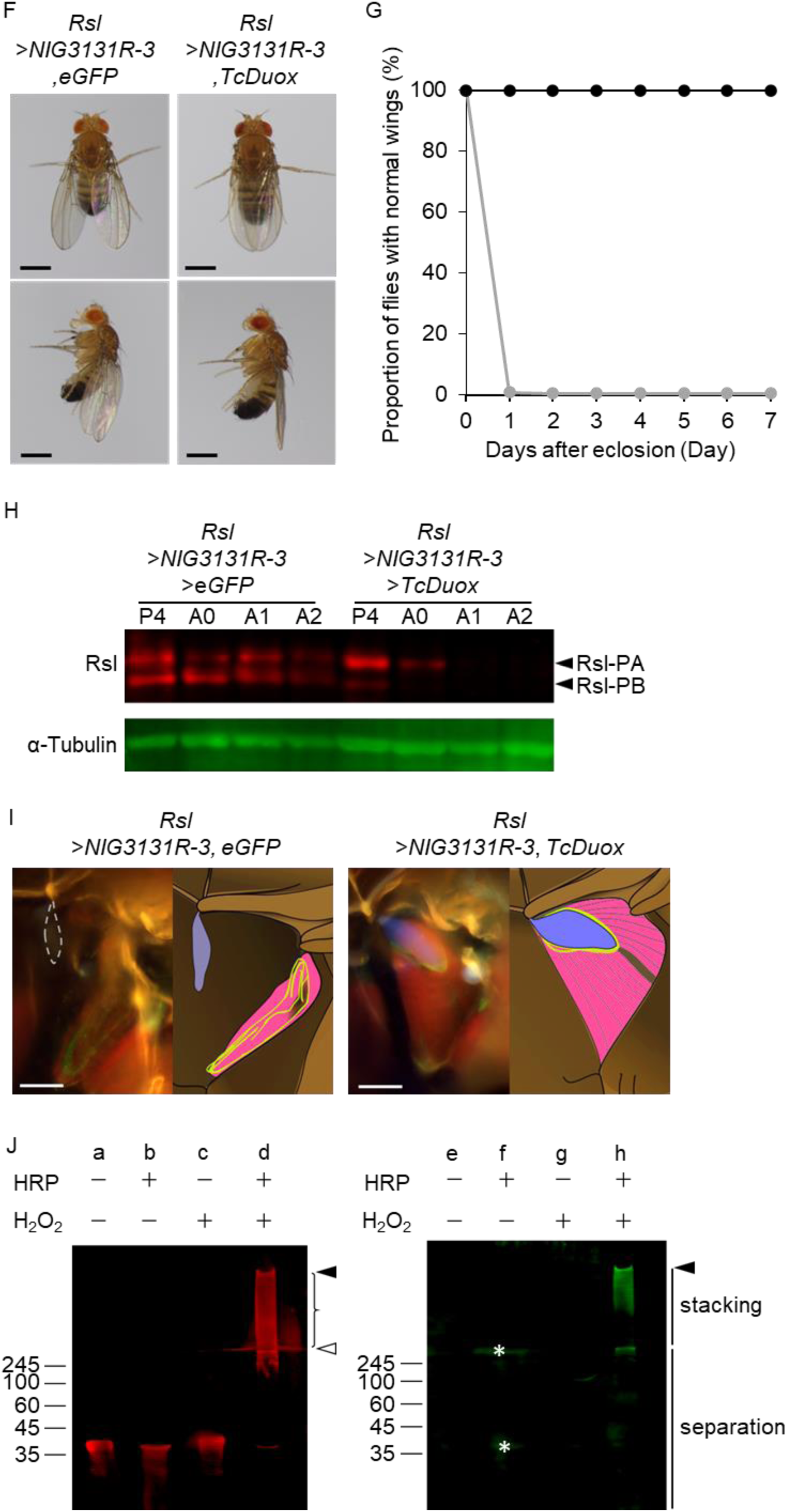
Analyses of *Duox* functions in Rsl polymerization. (A) The wing postures of the control (*Rsl>GFP-IR*) and *Duox* knockdown flies (*Rsl>NIG 3131R-3*, and *Rsl>Duox-RNAi_976–1145_*) are compared. Scale bar: 0.5 mm. (B) The temporal changes in the proportions of flies with normal wing posture after eclosion are shown. The flies used were the same as those in (A) (closed circles: *Rsl>GFP-RNAi*, grey closed circles: *Rsl>NIG3131R-3*, and gray open circles: *Rsl>Duox-RNAi_976–1145_*). (C) Western blot analyses of Rsl protein levels were performed using samples from day-4-APF pupae to day-2-after-eclosion adults. The left and right are flies of the control (*Rsl>GFP-RNAi*) and *Duox* knockdown (*Rsl>NIG3131R-3*), respectively. (D) The possible model of Duox-mediated Rsl polymerization (after Edens et. al., 2001^38^) (E) The inner structures of the wing hinge in the two control flies, *Rsl-Gal4/+* (panel a) and *NIG3131R-3/+* (panel b), are shown. The changes in the structure in the *Duox*-knockdown flies (*Rsl>NIG3131R-3*) are shown (panels c and d). Panel e on the right is the schematic of panels c and d. TP-1 and TP-2 muscles were stained with phalloidin. In *Duox*-knockdown flies, the dotted line (panels c and d) indicates the outline of the dityrosine signal corresponding to the weak blue fluorescence signal indicated by triangles in the photo taken with a longer exposure time. Scale bar: 50 µm. (F) Wing posture of the *Duox* knockdown fly (*Rsl>NIG3131R-3, eGFP*) and that of the *Duox* knockdown fly with the additional *TcDuox* expression (*Rsl> NIG3131R-3, TcDuox*). Scale bar: 0.5 mm. (G) Difference in the proportion changes of flies with normal wing posture was compared between *Rsl> NIG3131R-3, eGFP* (gray circle) and *Rsl>NIG3131R-3, TcDuox* (closed circle) flies. (H) Western blot analyses were performed to examine the levels of the Rsl protein from day-4-APF pupae to day-2-after-eclosion adults are shown. The left shows the *Duox*-knockdown individuals (*Rsl>NIG3131R-3, eGFP*) and the right shows the rescued individuals (*Rsl>NIG3131R-3, TcDuox*). (I) The photos and illustrations of the inner structure of the wing hinge in *Duox*-knockdown (left panels) and rescued flies (right panels) are shown. Scale bar: 50 µm (J) Western blot analyses of reRsl protein and dityrosine levels. reRsl was incubated in the presence (+) or absence (-) of 20 µg/µL horseradish peroxidase (HRP) and/or 0.3% hydrogen peroxide (H_2_O_2_) in 50 mM sodium phosphate buffer (pH 7.0) for 1 h at 30°C. Closed arrowheads show the signals stacking at the entry site of the concentration gels. The open arrowhead shows the signal stacking at the border between the concentration and separation gels. Curly bracket shows smear signals in the stacking gel.

To examine the cause of the downturned wing posture in *Duox* knockdown flies, we performed western blot analysis to determine whether Rsl molecules were polymerized. As shown in Figure 4C, in the *Duox* knockdown flies, the signal for Rsl can be seen at the same electrophoretic mobility of Rsl monomers (Figure 4C, right panel), suggesting that a portion of Rsl molecules remained unpolymerized. This is in contrast to the patterns of the control (Figure 4C, left panel) and wild-type flies (Figure 2B), in which the Rsl signals disappeared after eclosion. Another explanation for the detection of *Rsl* in the *Duox* knockdown flies could be the enhancement of *Rsl* expression after becoming adults. However, there were no marked differences in the patterns of *Rsl* expression between the control and the *Duox* knockdown flies (Figure S9), which does not negate the model of the Duox-mediated Rsl polymerization (Figure 4D).

We also observed the structure around the tergopleural tendon of the *Duox* knockdown flies (Figure 4E). In the control flies (panels a and b), the structures were almost the same as those of the wild type (Figure 3F), as can be judged from the presence of the blue fluorescence signal at the tergopleural tendon. In the *Duox* knockdown flies, the signal for dityrosine in the tergopleural tendon (indicated by the dotted line in panel c) was very weak. In the *Duox* knockdown flies, the TP-1 and TP-2 were completely detached from the tergopleural tendon with the signal of Dpy (panel d), as illustrated in panel e.

To rescue the phenotypes caused by *Duox* knockdown, we prepared a UAS-line for expressing of *T. castaneum Duox* (*TcDuox*) as a transgene with a base sequence that is not similar to that of *D. melanogaster Duox*. As shown in Figure 4F, *Duox* knockdown flies with additional *TcDuox* expression did not exhibit the downturned wing posture, even one week after eclosion (Figures 4F and 4G). The flies could move their wings normally and fly (our observation). In western blot analysis, the Rsl signal disappeared within two days after eclosion (Figure 4H, right panel), in contrast to the control flies in which the Rsl signal remained detectable even after two days (left panel). The anatomical structure around the tergopleural tendon was almost the same as that of the wild type (Figure 4I, right panel), which can be judged from the presence of the blue fluorescence signal in the tergopleural tendon, and also the connection between the tergopleural tendon and TP muscles. These findings indicate that the expression of *TcDuox* rescued the phenotypes of the *Duox* knockdown flies, and that *Duox* is the actual factor responsible for Rsl polymerization and the proper functioning of Rsl/resilin matrix in the maintenance of the structure around the wing hinges. We then examined biochemically the Duox-mediated Rsl polymerization through dityrosine formation. Here, commercially available peroxidase was used as the catalyst, instead of the peroxidase domain of Duox, because we were unable to synthesize the functional peroxidase domain of Duox successfully. In the western blot analysis of the truncated version of the recombinant Rsl (reRsl), which was synthesized in an *E. coli* system, reRsl migrated at an electrophoretic mobility corresponding to the predicted molecular mass of 35 k (Figure 4J, left panel, lane a). After treatment with either peroxidase or hydrogen peroxide (lanes b and c), reRsl can be seen at its original mobility. In contrast, in the presence of both peroxidase and hydrogen peroxide, the reRsl signal was shifted to an electrophoretic mobility corresponding to a much higher molecular mass (lane d). As indicated by closed arrowheads, the strong smear signal was seen from the entry site of the concentration gel. The shift of the Rsl signal to an electrophoretic mobility corresponding to high-molecular masses indicates that the Rsl molecules formed a high-molecular mass complex. In the western blot analysis with the antibody against dityrosine (right panel), signals are visible only in the sample treated with both peroxidase and hydrogen peroxide (lane h). The signal is strongest at the entry site of the concentration gel (closed arrowhead), indicating the presence of reRsl, which is highly polymerized by the formation of a large number of dityrosines.

### Both the expressions of *Rsl* and *Duox* are required for high jump performance

It has been proposed that the resilin matrix may enable high performance in jumping in several insects, such as the flea and locust^10, 21,41,42^. In this study, as a method to quantify the jump performance in *D. melanogaster*, we measured the jump distance using de-winged flies (Figure 5A). In the control flies, their jump distance was around 2 cm, but in the *Rsl* knockdown flies, their jump distance was shorter by around half of that of the control (Figure 5B). Also, in *Rsl^KO^*flies, the jump distance was almost half of that of the wild-type flies (Figure 5C, Extended data Video 4 and 5). The jump distance of the *Rsl^KO^* flies with *Rsl-GR* was almost the same as that of the wild-type flies (Figure 5C), indicating that the phenotype of shorter jump distance was rescued by *Rsl-GR*. In the rescue experiment using the Gal4/UAS system, we were unable to obtain data clearly showing the successful rescue with either *UAS-Rsl-RA* or *UAS-Rsl-RB*. Therefore, presently, we do not have information on the functional differences in jumping between Rsl-PA and Rsl-PB (not shown).

**Figure 5.**
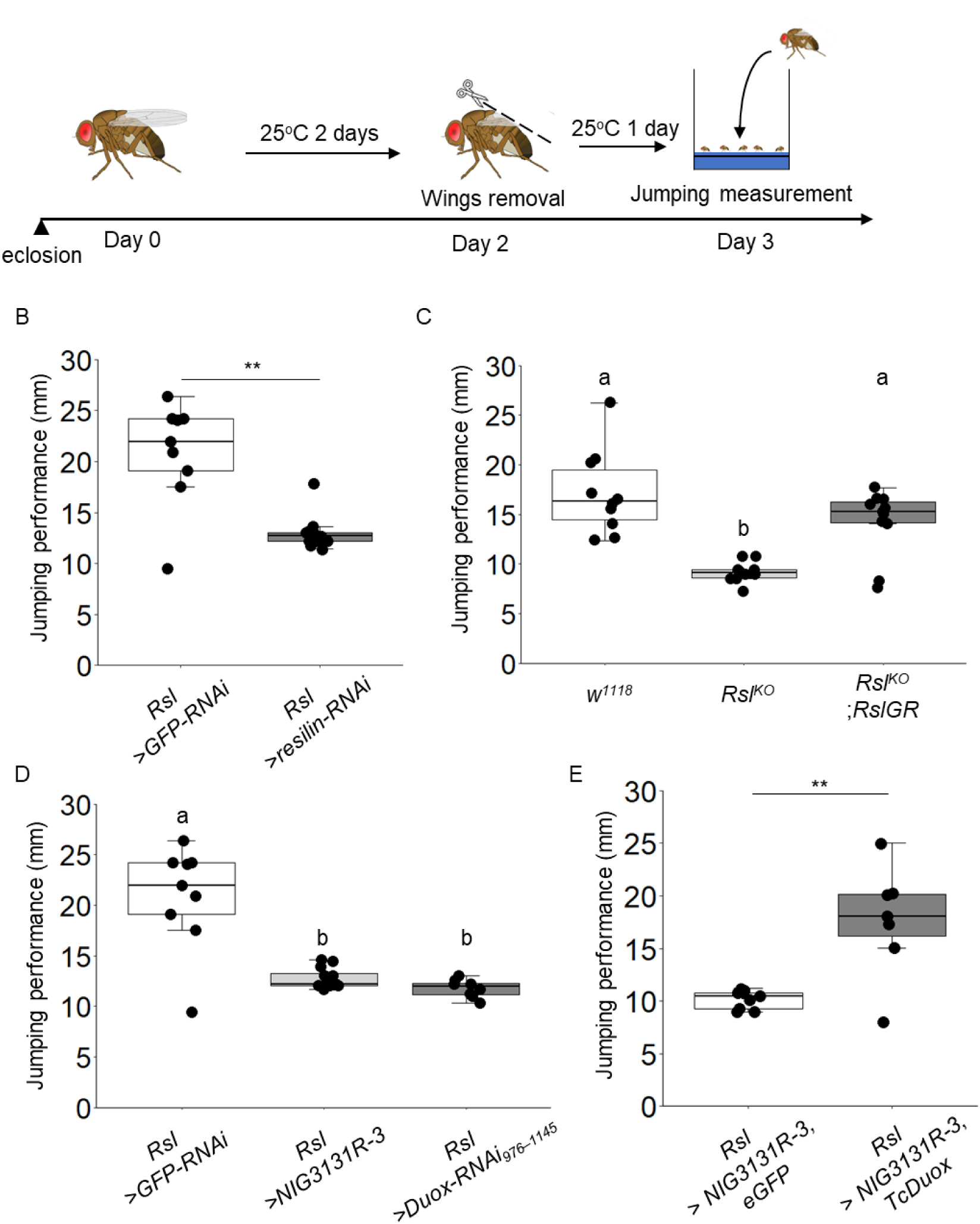
Contributions of *Rsl* and *Duox* to fly’s jumping. (A) Experimental scheme to measure jump distance of de-winged flies. (B) The jump distances between the control flies (*Rsl>GFP-RNAi*, white box) and *Rsl* knockdown flies (*Rsl>Rsl-RNAi*, gray box) are compared. Cross marks indicate the average. (Wilcoxon rank sum test, **: *P* < 0.01, *Rsl>GFP-RNAi*: n=8, *Rsl>Rsl-RNAi*: n=12). (C) The *Rsl^KO^* phenotype of jump distance was rescued with *Rsl-GR*. The flies used were the wild-type (*w^1118^*, white box), *Rsl^KO^* (*w^1118^*; *Rsl^KO^,* gray box), and *Rsl^KO^* with *Rsl-GR* (*w^1118^*; *Rsl^KO^*; *Rsl-GR*, dark gray box). Cross marks indicate the average (Wilcoxon rank sum test, Bonferroni correction, *w^1118^*: n=10, *Rsl^KO^*: n=10, *Rsl^KO^* with *Rsl-GR*: n=11). (D) Effect of *Duox* knockdown on jump performance was examined. The graph shows the jump distances of the control flies (*Rsl>GFP-RNAi,* white box), the *Duox* knockdown flies (*Rsl> NIG 3131R-3*, indicated by the light gray box, and *Rsl*>*Duox-RNAi_976–1145_* indicated by the gray box) (Wilcoxon rank sum test, Bonferroni correction, *Rsl>GFP-RNAi*: n=9, *Rsl> NIG 3131R-3*: n=12, *Rsl*>*Duox-RNAi_976–1145_*: n=8). (E) The phenotype of *Duox* knockdown flies was rescued by the additional expression of *TcDuox*. The graph shows the jump distances of the *Duox* knockdown flies (*Rsl> NIG3131R-3, eGFP,* white box) and the flies with *TcDuox* expression (*Rsl> NIG3131R-3, TcDuox*, gray box) (Wilcoxon rank sum test, **: *P* < 0.01, *Rsl> NIG3131R-3, eGFP*: n=9, *Rsl> NIG3131R-3, TcDuox*: n=7).

Here, in addition to *Rsl*, we examined the contribution of *Duox* to fly jumping. *Duox* was knocked down with either of *NIG3131R-3* or *UAS-Duox-RNAi_976-1145_*, and in both cases, the jump distances of the knockdown flies were around half of that of the controls (Figure 5D). To rescue this phenotype, *TcDuox* was additionally expressed in the *Duox* knockdown flies (*Rsl*>*NIG3131R-3*). With the expression of *TcDuox*, the jump distance was increased to around twofold that of the *Duox* knockdown flies (Figure 5E), indicating that *TcDuox* expression compensated for the reduction in the level of the endogenous *Duox* expression.

## Discussion

Duox is the oxidase that is widely conserved in metazoans, and is known as the thyroid oxidase in mammals^43,44^. There have been many studies on the diverse roles of Duox, including defense responses against intestinal wounds, immune responses against pathogens, the regulation of stem cell development/differentiation, and extracellular matrix formation^37–40,45–48^. In particular, in ecdysozoans such as insects and nematodes, it is considered that the Duox-mediated crosslinking through dityrosine formation is very important for the strengthening and stabilization of the extracellular matrix^37–39,49^. In this study, we showed that Rsl was polymerized by Duox, in which inter-Rsl molecule dityrosine formation may occur (Figures 4C, 4D, 4H, and 4J). In insects, this is the first study to identify the target protein of the Duox-mediated polymerization. The polymerization of Rsl may be used as a model to study the formation of the dityrosine matrix, which is distributed widely in metazoan taxa, including annelids and molluscs^50,51^.

Until this study, two reports on the function of the resilin-encoding gene in Drosophilae species had been published, both by Moussian’s group. Here, we were able to specify the internal structures related to the emergence of this phenotype (Figure 3F). As shown in Figure 3F, the tergopleural tendon was ruptured in *Rsl^KO^* flies. The direct flight muscles (TP-1 and TP-2) that are linked to the tergopleural tendon are required for the movement of the wings that are upturned, as has been demonstrated in a neurogenetic experiment^52^. Probably, in the *Rsl* knockdown and *Rsl^KO^* flies, the tergopleural tendon may have been damaged owing to the accumulation of stress from the wing movement, since the occurrence of this phenotype is time-dependent (Figures 3B and 3D). After the rupture of the tergopleural tendon, energy transfer from the muscle to the wings may no longer occur.

In the insect database of the full-length CDS or long-read RNA-seq results, we found two splicing isoforms of resilins with or without the chitin-binding domain (our observation), which indicates the possibility that the presence of these two isoforms is important. The localization of Rsl-PA::mCherry (with the chitin-binding domain) in the tergopleural tendon (Figure 2H) is actually similar to that observed in the distribution of chitin fibers that were visualized using the ChtVis-tdTomato chitin reporter (Figure 2I). This suggests that the chitin-binding domain of Rsl-PA is functional *in vivo*, which is consistent with the finding of an *in vitro* experiment using recombinant Rsl, showing the binding of the chitin-binding domain to chitin^23^. The binding of the Rsl molecule to chitin fibers seems to have a certain significance for the formation and proper functions of the tergopleural tendon. However, when we compared the extent of phenotypic rescue by overexpressions of *Rsl-RA* and *Rsl-RB* in *Rsl^KO^*flies, the downturned wing phenotypes were rescued, in almost the same proportion (Figure 3D). This suggests that the chitin-binding domain is not necessarily required for Rsl functions at the wing hinge. In the wing hinge, it is possible that other cuticle proteins with both tyrosine residues and the chitin-binding domain may compensate for the absence of the chitin-binding ability of Rsl-PB by mediating the linkage between chitin fibers and Rsl-PB.

The polymerization of elastic protein moieties often occurs to give the matrix high elasticity, as seen in other elastic materials, such as elastin and abductin^53–55^. *In vitro* experiments designed to polymerize recombinant Rsls support this theory^5,20,53^. In these studies, either peroxidase, photocatalysts, or chemical modifiers were used to artificially polymerize Rsl, and the resulting materials treated with these reagents exhibited high elasticity. The representative molecule forming the elastic matrix in mammals is elastin, which is referred to as the “molecular spring” because of its roles in restoring the structure and providing elasticity to tissues^53–55^. The tyrosine residues of Rsl may form intermolecular crosslinks, which may be responsible for the formation of the resilin matrix and its high resilience. According to the UniProt website (https://www.uniprot.org/uniprotkb/Q9V7U0/entry), the chitin-binding domain of the resilin model has a pLDDT score greater than 90, indicating high precision in the 3D model prediction. However, the scores for the N- and C-terminal repeats are estimated to be low (pLDDT score <70), which indicates that these regions could be intrinsically disordered proteins. This is consistent with previous predictions that the Rsl molecule has a random structure, lacking alpha helices or beta sheets^23,56^. However, low scores in AlphaFold’s predictions may simply reflect insufficient information about proteins possessing similar sequences that could serve as references for known protein structures during 3D modeling. At present, there is insufficient evidence to rule out the possibility that Rsl possesses a structure similar to that of the predicted model. To accurately understand the thermodynamic mechanism by which polymerized Rsl molecules exhibit high elasticity, it would be beneficial to confirm whether Rsl actually possesses a coil-like structure.

As shown in Figure 4A, the knockdown of *Duox* induced the same downturned wing posture as that of the *Rsl* mutant. This result can be explained by the failure of Rsl polymerization, as shown in the western blot in Figure 4C. Even in the adult stage, Rsl polypeptides remained detectable at an electrophoretic mobility corresponding to those of the Rsl monomers. *Duox* knockdown also induced the reduction in the intensity of dityrosine signal in the tergopleural tendon (Figure 4I), strongly supporting the idea that Duox is the factor for dityrosine-mediated Rsl polymerization. The reduction or loss of the elasticity of the resilin matrix at the tergopleural tendon may have occurred, finally leading to the rupture of the tergopleural tendon. It is also possible that the link between the tergopleural tendon and the muscle is lost, probably because of the weak bonding between them, as can be assumed from the detachment of the muscle from the tergopleural tendon (Figure 4E).

The model that Duox is the factor for Rsl polymerization was based on the studies of the nematode *Caenorhabditis elegans*, showing that the gene for Duox is involved in the dityrosine-mediated polymerization of collagens in the cuticle matrix^38,49^. It is possible that this process may be controlled by humoral factors, because it is stage-dependent (Figure 2B). It has been shown that the functions of Duox are regulated by the EF-hand motif present at the intracellular part of this protein^40,45^. We observed an increase in the cytosolic calcium level in the cells surrounding the tergopleural tendon in the late pupal stages (our observation). The next step is to clarify whether the increase in the cytoplasmic calcium level actually contributes to the activation of Duox and the subsequent dityrosine formation, followed by elucidation of the molecular mechanisms underlying the stage-dependent regulation of Rsl polymerization during the process of adult development.

Regarding jumping, both *Rsl^KO^* and *Duox* knockdown flies showed shortened jump distances, suggesting the loss of the polymerized Rsl may have been the cause of this phenotype (Figure 5). In a recent study on the resilin gene of the locust *S. gregaria*, the authors showed the lower jump performance of RNAi individuals (reduced jump velocity and irreversible reduction in the jump distance during repetitive jumps)^57^. It was also shown that the joints of the hind legs (between the femur and the tibia) were more easily broken in the RNAi individuals. In *D. melanogaster*, the resilin matrix is present on the ventral side of the leg joint (Figure 2); from the finding, we proposed a simple model on how the resilin matrix works in fly jumping (Figure S10). Before jumping, the resilin matrix becomes tense to store mechanical energy, and when the fly jumps, the stored energy may be liberated from the resilin matrix during transition to its relaxed state.

As shown in the observation of the *Rsl^KO^* flies, we found the wing structures still showing blue fluorescence (Figure S8). The lack of Rsl is also supported by the previous observation that there are no reporter signals at the proximal part of the wing^21^. In many reports, the term ‘resilin’ has been used to refer to the cuticular or other extracellular structures with blue fluorescence. However, the data from *Rsl^KO^*flies show the existence of the non-resilin matrix with dityrosine, indicating that the matrices positive for blue fluorescence do not necessarily contain Rsl molecules. It is possible that such the non-resilin matrix with blue fluorescence may have mechanical properties quite different from those of the resilin matrix, because blue fluorescence can be found in very hard parts such as the tip of the mandible of copepods or ragworm^58,59^. Future studies will reveal the mechanical characteristics or composition of the non-resilin matrix with blue fluorescence and the biological significance of their existence.

## Materials and Methods

### Fly stocks and maintenance

Fly stocks were reared at 25°C under a 12h light and dark cycle. They were fed the standard cornmeal–agar–glucose medium containing propionic acid and n-butyl p-hydroxybenzoate (7.21% cornmeal, 3.20% glucose, 8.01% yeast, and 0.64% agar supplemented with 9.9 mL/L butyl *p*-hydroxybenzoate and 3.0 mL/L propionic acid). *w^1118^* (our laboratory stock, Canton-S background) was used as the wild-type strain. *UAS-eGFP* (BL#1521), *UAS-GFP-RNAi* (BL#9331), *vas-int* (BL#40161), AttP2 (BL#36303), AttP5 (BL#34766), and *vas-Cas9* (BL#51323) were obtained from the Bloomington Drosophila Stock Center. *UAS-resilin-RNAi* (*KK106773*) in VIE260b (#101068) was obtained from the Vienna Drosophila Resource Center. *UAS-Duox-RNAi* (#3131R-3) and *y^2^cho^2^v^1^; Sco/CyO* (#TBX-0007) were obtained from Fly Stocks of the National Institute of Genetics. *UAS-Duox-RNAi* ^40^ was kindly provided by Dr. Won-Jae Lee (Seoul National University, Korea). All mutants and lines that were used for knockdown were crossed into the *w^1118^*control background for at least five generations.

### RNA isolation and reverse-transcription PCR

Total RNA was extracted from individuals at each growth stage using RNAiso Plus (D9108A, Takara, Japan). cDNA was synthesized from 1 μg of total RNA using Prime Script RT Reagent Kit with gDNA Eraser (DRR047A, Takara). cDNAs were amplified using Prime star Max (R045A, Takara). To obtain patterns of stage-specific gene expressions, a three-step thermal cycling profile was used as follows: 95 °C for 1 min, followed by 25 cycles at 98 °C for 10 s, 55 °C for 5 s, and 72 °C for 5 s. The primers used in the RT-PCR are listed in Supplementary Table 1.

### Generation of fly strains

*Rsl^KO^* was obtained by the CRSPR/Cas9-mediated deletion of the whole coding region with two gRNAs^60^. The gRNAs were expressed from a transgene with a ubiquitous *pol III* promoter derived from *Drosophila snRNA:U6:96Ab*^61^. For construction of DNA to generate flies expressing gRNAs, two primer sets for *Rsl-gRNA1* and *-gRNA2* (listed in Supplementary Table 2) were annealed, and the resulting DNA fragments were separately subcloned into *pBFv-U6.2* and *pBFv-U6.2B*, respectively, that were predigested with *Bbs*I. *pBFv-U6.2* with the sequence of *Rsl-gRNA1* was digested with *EcoR*I and *Not*I, and one of the resulting DNA fragments (with the sequence of *Rsl-gRNA1*) was ligated into one of the *EcoR*I and *Not*I fragment of *pBFv-U6.2B* (with the sequence of *Rsl-gRNA2*). The resulting vector with both the sequences for expressing *Rsl-gRNA1* and *-gRNA2* was introduced into AttP2 flies using a phiC31 integration system^60,61^. The flies with the transgene were crossed with *vas-Cas9* flies and the offspring were crossed with *y^2^cho^2^v^1^; Sco/CyO* virgins to establish lines with deletions. The genotypes of the established lines were checked by genomic PCR analysis using the primers listed in Supplementary Table 2.

For the construction of the plasmid vector for generating transgenic flies (*Rsl-Gal4, Rsl* genomic rescue (*Rsl*-GR)*, UAS*-*Rsl-RA*, and *UAS*-*Rsl-RB*), genomic DNA and cDNA obtained from *w^1118^* were used as templates of PCR. For the subcloning of the PCR fragments, an In-Fusion HD cloning kit (Z9648N, Takara) was used. To construct the vector of *Rsl-Gal4*, the 2.3 kbp PCR fragment for the *Rsl* promoter region amplified using the primer set (Supplementary Table 2) was subcloned into *pBPGw* (Addgene; https://www.addgene.org/) that was predigested with *Kpn*I and *Bgl*II. To generate flies with the transgene for the genomic rescue of *Rsl^KO^*, we prepared a 4.7 kb genomic fragment containing the *Rsl* CDS (from 1.0 kb upstream to 0.5 kb downstream of the transcription start and stop sites of *Rsl*) which was amplified by PCR using the primer set (Supplementary Table 2). The DNA fragment was subcloned into *pBFv-U6.2* predigested with *EcoR*I and *Not*I. For the construction of the plasmid vector to generate *UAS-Rsl-RA* and *UAS-Rsl-RB*, PCR fragments corresponding to the whole coding regions of *Rsl-RA* and *Rsl - RB,* respectively, were amplified using the primer set (Supplementary Table 2) with the cDNA. They were subcloned into *pUAST-attB* (laboratory stock) predigested with *Not*I and *Xba*I. For the construction of the vector for *UAS-TcDuox*, the DNA fragment corresponding to the whole coding region of the duox gene of the red flour beetle *Tribolium castaneum* was amplified by PCR using the *T. castaneum* cDNA and the primer set (Supplementary Table 2) (*T. castaneum* was a kind gift from Dr. Toru Togawa, Nihon University, Japan). For the construction of *UAS-Rsl-RA::mCherry* and *UAS-Rsl-RB::mCherry*, the whole coding region of each isoform was amplified by PCR using the primer set (Supplementary Table 2) with cDNA from iso-1 (BL2057) as the template. A fragment including the sequence of mCherry with an N-terminal flexible 4 × GlyGlySer linker was PCR-amplified from *UASp-Elyssim-mCherry*^62^ using the primer set (Supplementary Table 2). The two DNA fragments were subcloned into *EcoR*I fragment of *pUAST-attB* by the SLiCE method (Motohashi, 2015). Each vector was introduced into flies with AttP5 (*UAS-Rsl-RA, UAS-Rsl - RB,* and *UAS*-*Tcduox*) or AttP2 (*Rsl-Gal4*, *Rsl-GR, UAS-Rsl-RA::mCherry*, and *UAS-Rsl-RB::mCherry*) using a phiC31-based integration system^63,64^.

### Generation of antibody

A part of Rsl (Lys478-Gly590) was chosen as the antigen to raise the antibody against Rsl. In this study, pET-32 (Novagen, USA) was used to express the Rsl antigen as a fusion protein with the N-terminally located thioredoxin. The DNA fragment corresponding to the antigen was amplified with the primer set (Supplementary Table 1) using the cDNA of *w^1118^*. The PCR product was digested with *Xho*I and *Bgl*II, and ligated into pET-32 predigested with the same enzyme set. The vector constructed was used to transform the *E. coli* BL21 strain. After the IPTG induction of the recombinant protein expression (1 mM, 4 h of culture at 37°C), the *E. coli* cells were harvested by centrifugation, and ruptured by sonication in 50 mM Tris-HCl (pH 7.5) containing 150 mM sodium chloride. From the extract, the recombinant protein was purified by Ni-NTA affinity chromatography using Ni-NTA Agarose (30210, Qiagen, Germany). The N-terminally located thioredoxin was removed by thrombin treatment () to break the peptide bond between the thioredoxin and the Rsl antigen, and the Rsl antigen was used for the immunization of a rabbit. The serum obtained from the rabbit was used as the anti-Rsl antibody.

### Western blot analysis of fly samples

Five individuals at different stages (white pupae, pupae, and adults) were homogenized in 400 µL of 2×SDS-PAGE sample buffer (12.5 mM Tris at pH 6.8, 20% glycerol, 4% SDS, 2% 2-mercaptoethanol, and 0.001% bromophenol blue) and boiled for 10 min at 100°C. The proteins in a 5 µL aliquot were separated by SDS-PAGE using 12.5% polyacrylamide gel and transferred to PVDF membranes (IPFL00010, Merck KGaA, Germany). After blocking using 5% skim milk with TBST (0.1% Tween20), the membranes were incubated with the rabbit anti-Rsl antibody (1: 3000 dilution) and mouse anti-α - Tubulin antibody [DM1A] (ab7291, Abcam, UK) (1: 5000 dilution) in Can Get signal solution 1 (NKB-201, Toyobo, Japan) at 4°C overnight. For the secondary antibody, IRDye-conjugated anti-rabbit and anti-mouse IgGs (925-68071 and 925-32210, respectively, LI-COR, USA) were used with Can Get signal solution 2 (NKB-301, TOYOBO, Japan) (1: 20000 dilution, for 2 h at 25°C). For luminescence imaging, an Odyssey DLx imaging system (LI-COR, USA) was used.

### Measurement of jump distance

One day before the experiments, the flies were de-winged under mild CO_2_ anesthesia. Jump distance was measured using the equipmentwhich was set in a room kept as 25°C. At the bottom of the glass beaker, 2% agarose was placed. After hardened, the agarose solution, a mesh for scaling was put on the agarose gel, and the surface was covered with additional 2% agarose. Five flies per group were housed in the equipment and their jumping were tracked for 5 minutes at 480 fps with Tough TG-5 (4545350-051112, OLYMPUS, Japan) and the jump distances were measured on the basis of intervals of the mesh. The longest jump distance in each group was used as the representative value. The experiments were performed at least eight times for each genotype and sex.

### Crosslinking of Rsl

Horseradish peroxidase was used for the crosslinking reaction of recombinant Rsl^21,65^. 100 µg of the recombinant Rsl (CSB-EP892693DLU, WUHAN HUAMEI BIOTECH, USA) was dissolved with 100 µl of 50 mM potassium phosphate buffer (pH7.0). Horseradish peroxidase (P-8375, Sigma, USA) was dissolved with 0.1 M potassium phosphate buffer (pH 6.0) to a final concentration of 20 µg/µL. 1 µL of the recombinant Rsl solution was mixed with 5 µL of 50 mM potassium phosphate buffer (pH 7.0). Then, 1 µL of horseradish peroxidase solution and 3 µL of 0.3% hydrogen peroxide (18084-00, KANTO CHEMICAL, Japan) were added, and the mixture was incubated at 30°C for 1 h. After incubation, the reaction mixture was mixed with 10 µL 2×SDS-sample buffer and boiled for 10 min at 100°C. Western blot analyses of Rsl and dityrosine were performed using the rabbit anti-Rsl antibody (this study) and the mouse monoclonal anti-dityrosine antibody (MDT-020P, JaICA, Japan). The experimental procedures of western blot analysis were the same as those described in section 2.5.

### Histology and microscopy

To observe the internal structure of the wing hinge, the adult thorax was divided into two parts (left and right) along the midline. Then the body parts around the wing hinge were obtained by trimming with microscissors. The tissues were fixed using 4% paraformaldehyde with PBS (pH 7.0). After blocking using 5% BSA (A7906, Merck KGaA) with PBST (0.05% Tween20), these samples were incubated in phaloidin-iFluor488 (CAY20549, Cayman Chemical, USA) with PBST (1:500) at 25°C overnight. An OLYMPUS BX60 (OLYMPUS, Japan) equipped with an WRAYCAM-VEX830 (WRAYMER, Japan) was used for whole-body or dityrosine, mCherry, and YFP signal detection. An OLYMPUS SZX7 (OLYMPUS, Japan) equipped with an OLYMPUS DP71 (OLYMPUS, Japan) was used for observing the phenotypes of wing posture.

### Quantification of biased Rsl::mCherry distribution

The standard deviation (std) and the mean intensity of the mCherry signal in the resilin matrix were quantified using ImageJ. The std/mean was used as the index of the biased Rsl::mCherry distribution^66^.

### Statistical analyses

Statistical analyses were performed using R version 3.6.0 (https://www.r-project.org/). The Wilcoxon rank sum test was used to compare the jump distances. The student t-test was used to compare the intensities of mCherry signals in the resilin matrix in the wing hinges. The Steel–Dwass test was used to compare the expression levels of *Rsl*, *Duox,* and *TcDuox*.

## Supporting information

w1118 wing movement

KO wing movement

Rescue wing movement

w1118 jump

KO jump

## Acknowledgements

*UAS-Duox-RNAi_976–1145_* was a kind gift of Dr. Won-Jae Lee (Seoul National University, Korea). *T. castaneum* was a kind gift from Dr. Toru Togawa (Nihon University, Japan). We are grateful to Mr. Kensuke Aizawa for his contributions to our preliminary experiments in the early period. We thank the National Institute of Genetics, the University of Indiana. Bloomington *Drosophila* Stock Center and the Vienna *Drosophila* Resource Center for providing fly lines. This work is supported by JSPS KAKENHI Grant Number JP23H02225 and 21J21904 and the Sasakawa Scientific Research Grant from The Japan Science Society.

## Author contributions

M.H. and T.Asa. initiated, designed the experiments, and developed the project. M.H. performed all the experiments, analyzed the data, and prepared the figures. Y.E. and Y.K. generated *UASt-resilin-RA::mCherry* and *UASt-resilin-RB::mCherry*. M.H. and T.Asa. wrote and edited the manuscript. H.O.I., T.S., T.A. and T.Asa. reviewed the manuscript.

