## Supplemental information for "*Resilin,* the gene for the molecular spring: its roles in flight and jumping of *Drosophila*"

Supplementary information

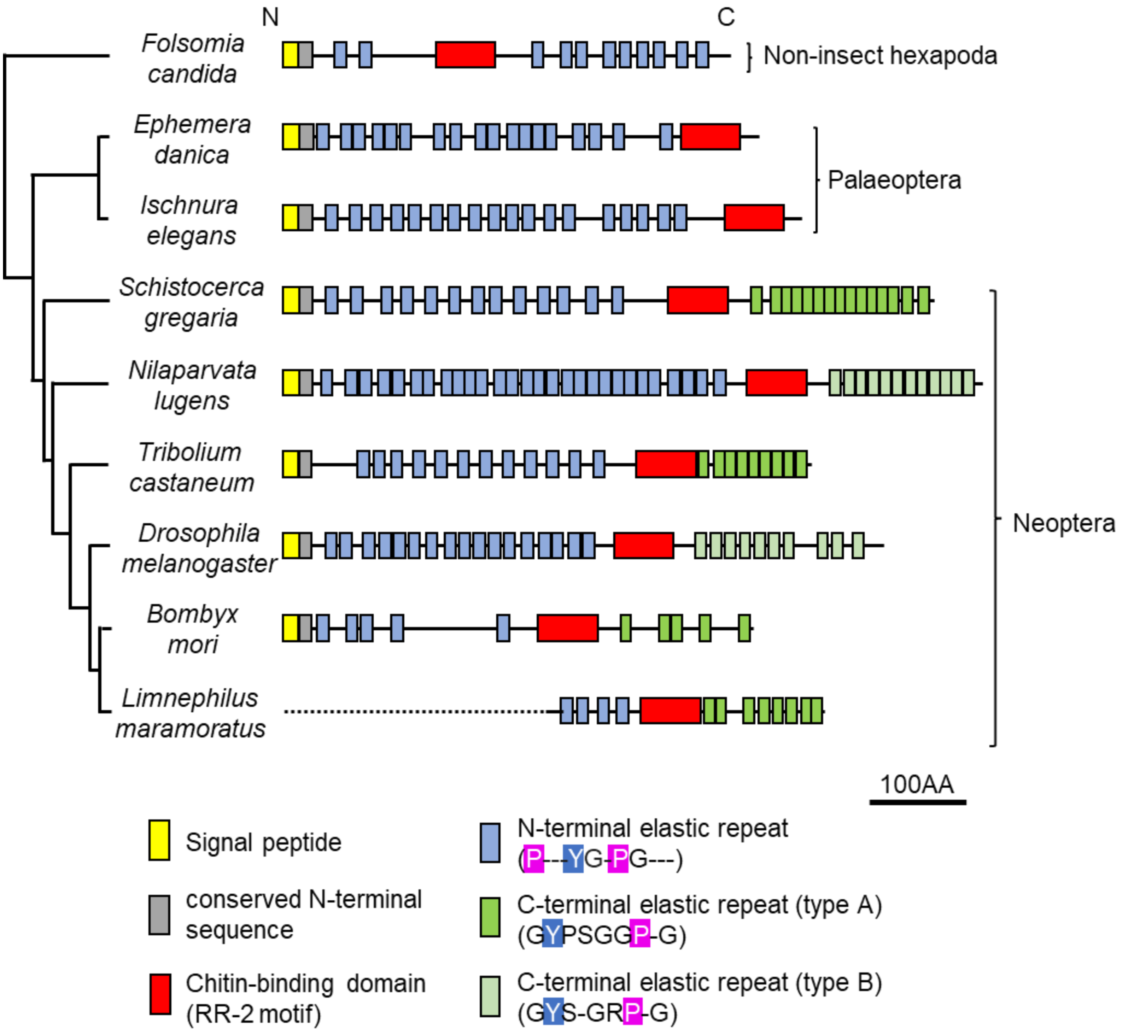


Figure S1: The domain/consensus organizations of resilins from Hexapoda were compared. The sequences used to construct the schematic structures are listed in Supplementary Table 3. The structure of this phylogenetic tree is based on the description in referece#67.

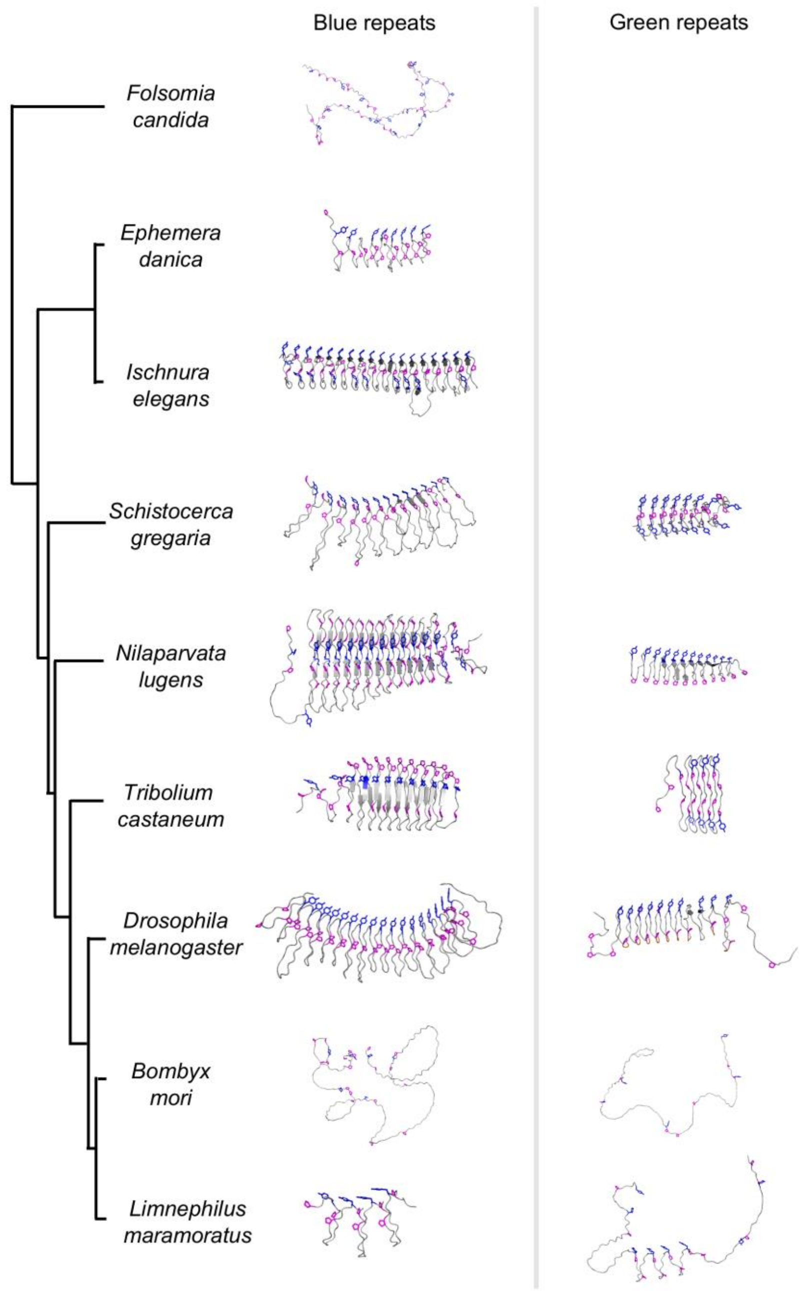


Figure S2: 3D models for elastic repeats of resilins from hexapods

For prediction in the AlphaFold server (https://alphafoldserver.com), the sequences listed in Supplementary Table 3 were used. Structural visualization was performed using PyMOL (The PyMOL Molecular Graphics System 2.5.4, Schrödinger, LLC).

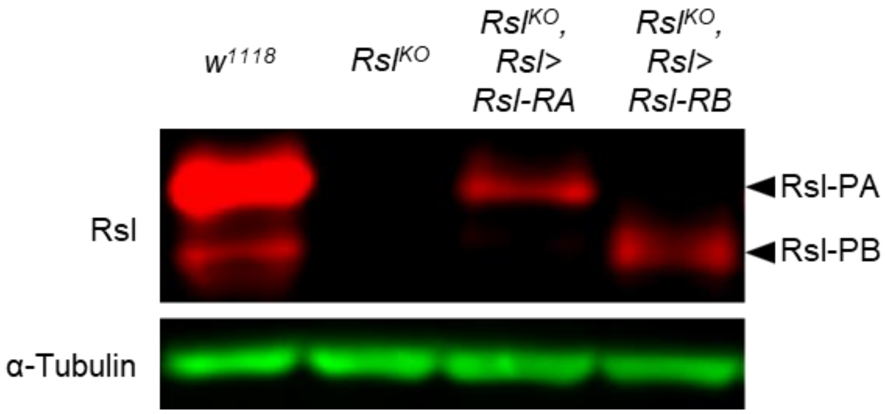


Figure S3: Western blot analysis of Rsl.

From the left, samples of $w^{1118}$, $Rsl^{KO}$, $Rsl^{KO}$ expressing *Rsl-RA* ($w^{1118}$; $Rsl^{KO}$, *UAS-Rsl-RA; Rsl-Gal4*), and $Rsl^{KO}$ expressing *Rsl-RB* ($w^{1118}$; $Rsl^{KO}$, *UAS-Rsl-RB; Rsl-Gal4*) were used. The samples were prepared with day-4-APF pupae. The experimental conditions are described in Materials and Methods.

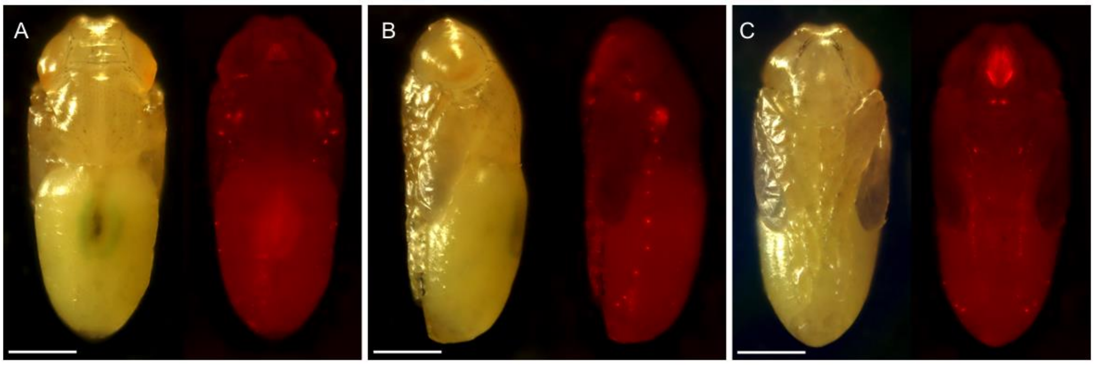


Figure S4: The pattern of Rsl::mCherry signals was expressed with *Rsl-Gal4* on the second chromosome. A –C are the day-4-APF pupae viewed from dorsal, lateral, and ventral directions, respectively. Scale bar: 500 µm.

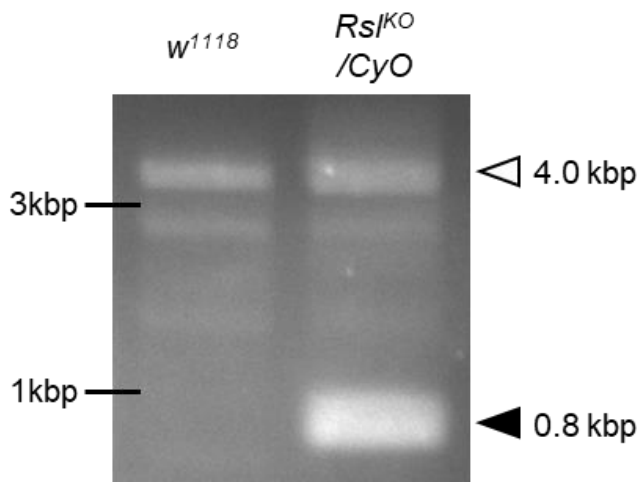


Figure S5: Genotyping of *$Rsl^{KO}$*. In genomic PCR with primers (Supplementary Table 2), two main bands are detected. The 4.0 kb and 0.8 kb fragments represent the *$w^{1118}$* (control) and *$Rsl^{KO}$* alleles, respectively.

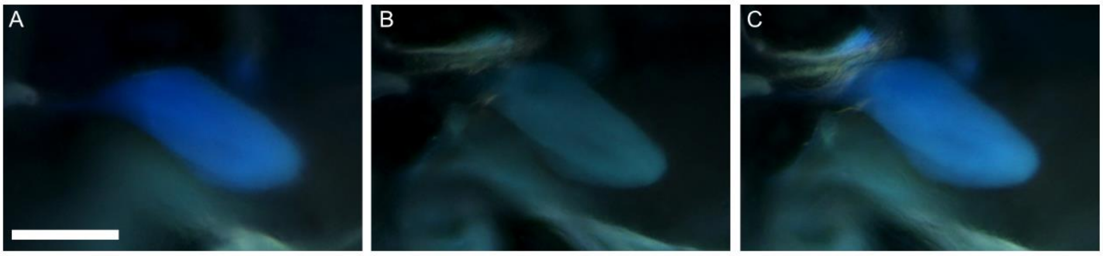


Figure S6: The resilin matrix in the hinge of wings in PBS (A), acidic solution (B, pH 2.0), and alkaline solution (C, pH 12.0). The blue fluorescence of dityrosine was drastically diminished in acidic pH but increased in alkaline pH under our experimental conditions.

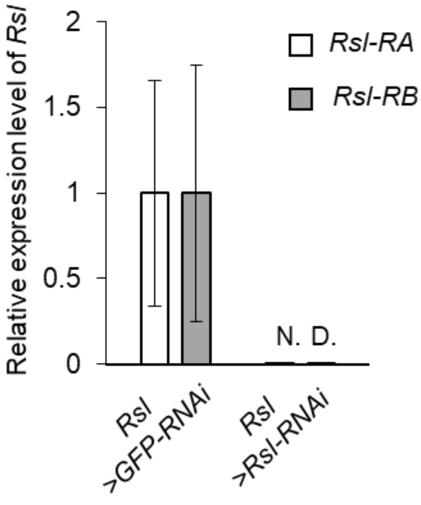


Figure S7: Relative expression levels of *Rsl-RA* and *Rsl-RB*.

Total RNA was extracted using RNAiso Plus. cDNA was synthesized from 1 μg of total RNA using Prime Script RT Reagent Kit with gDNA Eraser (DRR047A, Takara). RT-qPCR was performed using THUNDERBIRD SYBR qPCR Mix (QPS-201, TOYOBO, Japan) with LightCycler96 (Roche, Basel, Switzerland). A two-step thermal cycling profile was used as follows: 95°C for 1 min, followed by 45 cycles at 95°C for 15 s and 60°C for 45 s. Data were normalized with the value obtained for the ribosomal protein gene *Rp49*. In RT-qPCR, the same primers as those used in the RT-PCR (section 2.2) were used.

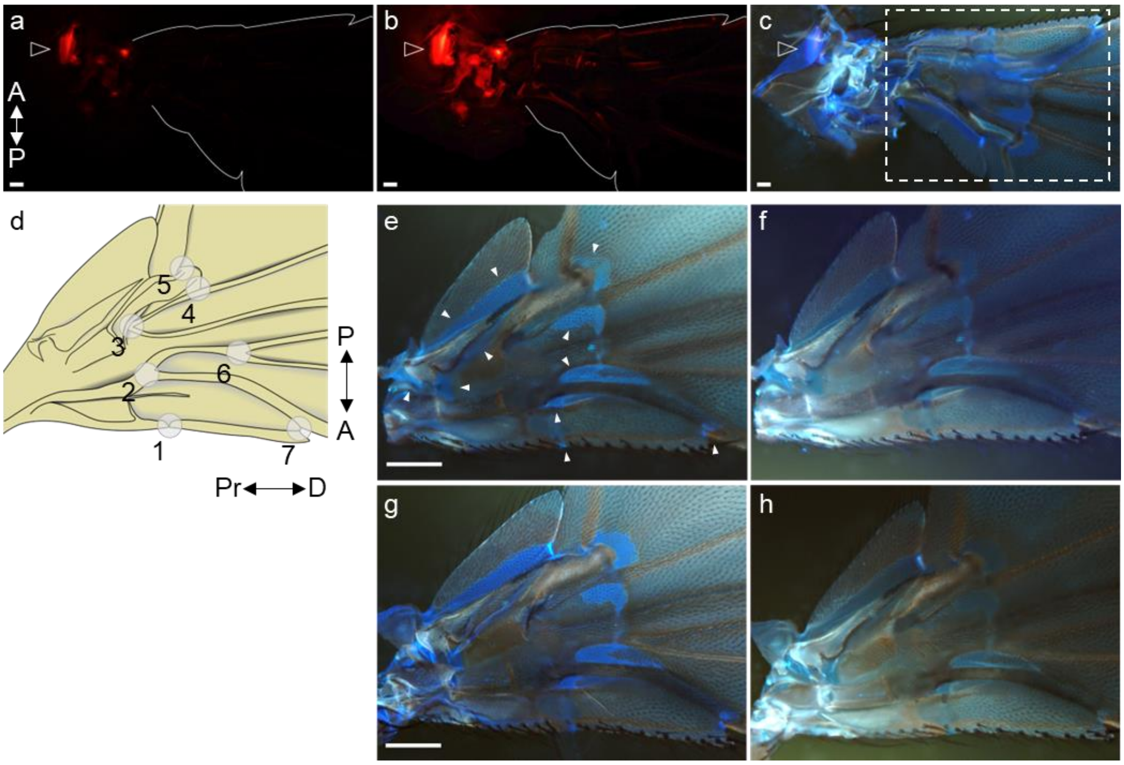


Figure S8: (a) The Rsl-RA::mCherry signal is seen at the tergopleural tendon (arrowhead). (b) Even in the photo, the brightness of which was increased, almost no signal was seen at the wing area indicated with the outline. (c) Under UV radiation, blue fluorescence signal can be seen at the wing. (d) A schematic showing landmarks in the wing (circles with numbers)[36]. (e and f) Photos in panels, e and f, show the wings of *w*$^{1118}$ immersed in PBSTs (pH 7.0 and pH 2.0, respectively). (g and h) Photos in panels, g and h, show the wings of *Rsl*$^{KO}$ immersed in PBSTs (pH 7.0 and pH 2.0, respectively). All photos were taken from the dorsal side (a, b, and c are the photos of right wing, and e, f, g, and h are the photos of left wings). White arrowheads indicate the dityrosine signals. Scale Bar: 100 µm

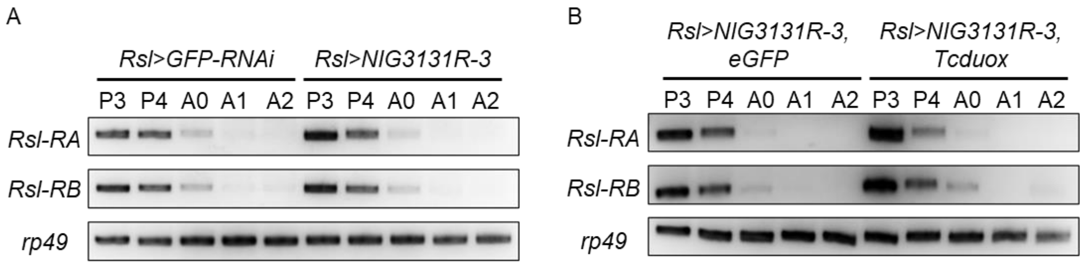


Figure S9: RT-PCR of each splicing isoform in *Rsl* from day-3-APF pupae to day-2-after-eclosion adults. (A) Control (*Rsl*>*GFP-RNAi*) and *Duox* knockdown (*Rsl*>*NIG3131R-3*). (B) *Duox* knockdown (*Rsl*>*NIG3131R-3*, *eGFP*) and knockdown rescued flies (*Rsl*> *NIG3131R-3*, *TcDuox*).

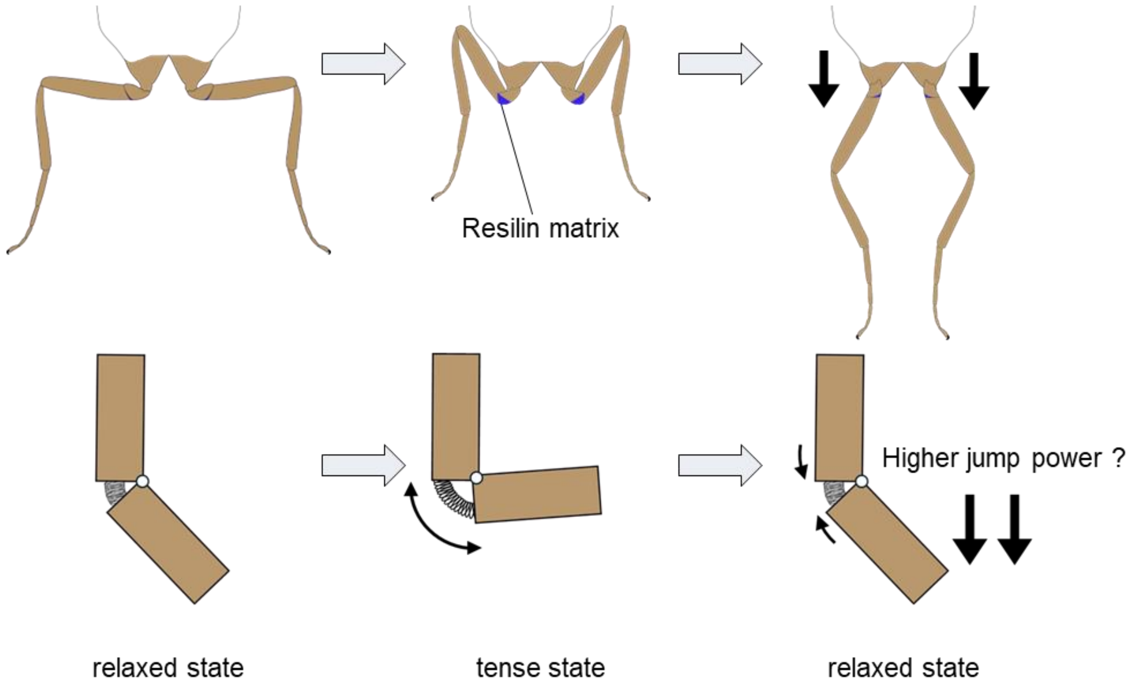


Figure S10: Model of Rsl functions in jumping. For details, see the text.

Supplementary Table 1: List of primers used for RT-PCR and qRT-PCR

| Supplementary Table 1 | |
|---|---|
| Primer Name | Sequence (5’ to 3’) |
| rp49 for | AAGATCGTGAAGAAGCGCAC |
| rp49 rev | TGTGCACCAGGAACTTCTTG |
| qPCR Resil Aisoform Fw | CGGAAGGAAGCAAATTGTGGAG |
| qPCR Resil Bisoform Fw | TGACTACGATAACGATATTGTGGAG |
| qPCR Resil Rv | TTTCCGTTGCCATTTCCATTGC |
| dDuox qPCR 4 Fw | AACAACAATGCTGTACATCTG |
| dDuox qPCR 4 Rv | GTATGAGTGCTTCTTTTGCAC |
| RT-PCR Tcduox Fw | CAACGCTATGACGGATGGTTCAAC |
| RT-PCR Tcduox Rv | TGCCCCGCTAGCATGTACAC |

Supplementary Table 2: List of primers used for the construction of vector plasmid

| Supplementary Table 2 | |
|---|---|
| Primer Name | Sequence (5’ to 3’) |
| *resilin* 5UTR _Forward | CTTCGATCTCCAGATGCGTCTGAT |
| *resilin 5*UTR _Reverse | AAACATCAGACGCATCTGGAGATC |
| *resilin 3*UTR _Forward | CTTCGTAGGGGACATAGTTCTAGA |
| *resilin* 3UTR _Reverse | AAACTCTAGAACTATGTCCCCTAC |
| resilinKO genotyping F | TGATCCATAATCCGCGAATGGCG |
| resilinKO genotyping R | GTAGCCCATGATCCAAAATACCG |
| resilin-Gal4 infusionF | ACGGCCGGCCAGATCTTTATATCCTGTACTTCTTGAGC |
| resilin-Gal4 infusionR | GGGATCCCGGATCTGCTTATTCCACTTGTATCTAAAGC |
| resil-rescue pBF infusionF | GCTTGATATCGAATT GTCCTAAGGCGATATAGACG |
| resil-rescue pBF infusionR | AGTGCATGCGCGGCC TACTAATATAAGTCGAACGG |
| resil UAS-exF | GCCGCGGCCGCGGAAATATGTTCAAGTTACTCGGCT |
| resil UAS-exR | CGGTCTAGACTAGTACCGATAACCGCTGCCATCGTT |
| Tcduox for NotI | CCGGCGGCCGCTCGAACATGGTTTCTTTAACATCTCTT |
| Tcduox rev XbaI | GAATCTAGACTAGCCGAAGTTTTCGAAGTGGTGGATGAAG |
| UASt-resilin::mCherry insert F | AGGGAATTGGGAATTCATGTTCAAGTTACTC |
| UASt-resilin::mCherry insert R | CGGATCCACCGCTGTACCGATAACC |
| linker-6xHis F | AGCGGTGGATCCGGCGGT |
| linker-6xHis R | GATCCTCTAGAGGTACCTTACTTGTACAGCTCGTC |

Supplementary Table 3: Accession numbers of Resilin in Hexapods used in this study.

| Species | Accession number | Source |
|---|---|---|
| *Folsomia candida* | XP_021963881.1 | NCBI |
| *Ephemera danica* | KAF4520732 | NCBI |
| *Ischnura elegans* | XP_046400398.1 | NCBI |
| *Schistocerca gregaria* | XP_049832388 | NCBI |
| *Nilaparvata lugens* | XP_022202958 | NCBI |
| *Tribolium castaneum* | BET60450 | NCBI |
| *Bombyx mori* | XP_012546125 | NCBI |
| *Limnephilus maramoratus* | ENSLMMT00005004393.1 | Ensembl Metazoa |
